# GPR68, a proton-sensing GPCR, mediates acid-induced visceral nociception

**DOI:** 10.1101/2025.06.13.659465

**Authors:** Luke W Paine, Rohit Gupta, James P Higham, Javier Aguilera-Lizarraga, Anne Ritoux, Thomas Pritchard, Samuel Nicholson, James R F Hockley, Tim Raine, Martin Hausmann, Kyle Bednar, Gerhard Rogler, Fraser Welsh, Ewan St John Smith, David C Bulmer

**Affiliations:** Department of Pharmacology, University of Cambridge, Tennis Court Road, Cambridge, CB2 1PD; Department of Gastroenterology, Addenbrookes Hospital, Cambridge University Teaching Hospitals, Cambridge, United Kingdom; Department of Gastroenterology and Hepatology, USZ, University of Zurich, Zurich, Switzerland; Respiratory & Immunology, BioPharmaceuticals R&D, AstraZeneca, Gaithersberg, USA; Neuroscience, BioPharmaceuticals R&D, AstraZeneca, Cambridge, UK

**Keywords:** visceral hypersensitivity, pain, nociceptors, colitis

## Abstract

**Background & Aims:** Localised acidification from immune cell infiltration and heightened glycolysis contributes to colitis pathology by activating acid-sensing receptors such as GPR68, a proton-sensing GPCR expressed on immune and stromal cells. Single-cell RNAseq analysis revealed GPR68 is also expressed in colonic sensory neurons, prompting us to investigate its role in acid-induced colonic nociception.

**Methods:** Expression of GPR68 in colonic nociceptors and tissue from people with colitis was confirmed by *in silico* analysis of our RNAseq databases. Its contribution to disease activity was assessed using the acute dextran sulphate sodium (DSS) model of colitis. Acid-evoked sensory signalling was evaluated via colonic afferent recordings and Ca²⁺ imaging in DRG neurons from wild-type and GPR68^-/-^ mice, supported by pharmacological studies using Ogerin (a GPR68 positive allosteric modulator) and Ogremorphin (a GPR68 antagonist).

**Results:** RNAseq analysis showed GPR68 is robustly expressed in *Trpv1*⁺ colonic nociceptors and upregulated in tissue from people with inflammatory bowel disease, consistent with reduced disease activity in DSS-treated GPR68^-/-^ mice. Genetic deletion of GPR68 abolished colonic afferent responses to acid, which were also attenuated by Ogremorphin and enhanced by Ogerin. In Ca²⁺-free buffer, DRG neurons from GPR68^-/-^ mice or those pre-treated with Ogremorphin showed significantly reduced acid-evoked intracellular Ca²⁺ responses. By contrast the colonic afferent and DRG Ca^2+^ response (in Ca^2+^-containing buffer) to capsaicin was comparable between tissue from wild-type and GPR68^-/-^ mice highlighting the involvement of divergent proton-dependent cellular signalling cascades.

**Conclusions:** These findings identify GPR68 as a key mediator of acid-induced colonic nociception and highlight its potential as a therapeutic target for the treatment of pain in colitis.

**Synopsis:** We demonstrate that the proton-sensing receptor GPR68 mediates acid-induced signalling in colonic sensory neurons. Genetic and pharmacological modulation confirms GPR68’s role in visceral nociception and supports its potential as a therapeutic target for pain management in colitis.

## 1. Introduction

Inflammatory bowel disease (IBD) is characterised by chronic inflammation of the gastrointestinal (GI) tract and is accompanied by acidification of the gut lumen and mucosa (1-4) with studies reporting colonic pH values ranging from pH 6.1-7.5 in healthy controls, 4.8-7.3 in ulcerative colitis (UC) and 5.3-6.5 in Crohn’s disease (CD) (1, 2), consistent with elevated faecal lactate levels (5) and reduced faecal pH (6). Local tissue acidosis is linked to IBD pathogenesis and can be attributed to the increased glycolytic activity, alongside reduced tissue perfusion and hypoxia (7), which occurs in response to the influx of immune cells, tissue inflammation and oedema (8). Furthermore, acidity drives pro-inflammatory gene expression in infiltrating immune cells through multiple acid-sensing mechanisms (9, 10) with emerging evidence suggesting that the upregulation and increased activity of proton-sensing G protein-coupled receptors (GPCRs), such as GPR68 contribute to these changes in immune cell behaviour and ultimately the pathogenesis of IBD (7).

The orphan proton-sensing GPCRs, GPR4, GPR65 (TDAG8), GPR68 (OGR1) and GPR132 (G2A), all have recognised roles in colitis pathophysiology (7, 11) and other inflammatory and neuropathic pain conditions (12-15). Although often classified with these receptors, GPR132 differs functionally and structurally, showing weak proton sensitivity and being primarily activated by oxidised fatty acids (16). Upon extracellular acidification, protonation of histidine residues in the extracellular domain of GPR4, GPR65, and GPR68 initiates receptor activation, a conserved feature among these core proton sensors (17). Recent computational and *in vitro* studies have expanded this model, implicating multiple cooperative interactions between histidine and acidic residues, across the extracellular and transmembrane domains, as critical for proton-sensing and conformational activation (18-20).

Intracellularly, GPR4 and GPR65 primarily couple to Gα_s_ proteins, whereas GPR68 preferentially couples to Gα_q/11_ proteins triggering phospholipase C (PLC) activation, inositol triphosphate (IP_3_) production and subsequent Ca^2+^ release from intracellular stores, which can lead to pro-inflammatory signalling cascades (17). Consistent with this, GPR68 activation perpetuates intestinal inflammation, and its genetic deletion confers resistance to fibrosis (21) and gut inflammation in IL-10^-/-^ spontaneous colitis models (22). Additionally, pharmacological inhibition of GPR68 decreases the severity of both acute and chronic DSS-induced colitis (23). GPR68 signalling also drives pro-inflammatory gene expression in immune and stromal cells, amplifying gut inflammation, and its expression is upregulated in colonic tissue during hypoxia (24-26). Consequently, GPR68 has emerged as a promising therapeutic target for inflammation in IBD.

In keeping with the underlying inflammation and tissue damage, abdominal pain is a leading cause of morbidity in colitis and its treatment represents a significant unmet clinical need facilitated by the contraindicated use of opioid and non-steroidal anti-inflammatory drug (NSAID)-based pain killers for use in people with IBD. Consequently, there is an increasing call from patient advocacy groups for the development of treatment strategies that can simultaneously address both inflammation and pain in IBD.

Given that acid-sensing is a conserved function of pain sensing neurons across species and a defining feature of polymodal nociceptors, facilitated by their expression of acid-sensitive ion channels and GPCRs (27), we sought to understand whether proton-sensing GPCRs such as GPR68 might also be responsible for the activation of colonic nociceptors by acid, and as such contribute to the activation of both nerve endings and immune cells following acidification of the bowel in individuals with colitis. GPR68 has previously been implicated in chemotherapy-induced neuropathy (28), and its heightened expression in sensory neurons (29) and GI tissue (22) compared to other proton-sensing GPCRs, supports a possible pro-nociceptive role for GPR68. Here, using complementary genetic and pharmacological approaches, we assess the contribution of GPR68 to acid-evoked sensory neuron and colonic afferent activation highlighting its potential as a therapeutic target for the development of novel visceral analgesics.

## 2. Methods

### 2.1. Ethical Approval

All experimental procedures performed on animals were conducted in compliance with the Animals (Scientific Procedures) Act 1986 Amendment Regulations 2012 under Project License PP5814995 granted to E. St. J. Smith by the United Kingdom Home Office with approval from the University of Cambridge Animal Welfare Ethical Review Body. In addition, animals were euthanised by rising concentration of 100% CO_2_ followed by exsanguination in accordance with Schedule 1 of the Animals (Scientific Procedures) Act 1986 Amendment Regulations 2012.

### 2.2. Animals

Adult C57BL/6J mice (8-16 weeks) were obtained from Charles River (Cambridge, United Kingdom; RRID: IMSR_JAX: 000664). GPR68 knockout (GPR68^-/-^) mice (MGI ID: 5776516) were kindly gifted by Martin Hausmann and Gerhard Rogler (24) and bred in Cambridge with C57BL/6J mice to generate wild-type (GPR68^+/+^) and heterozygous (GPR68^+/-^) littermates. Mice were housed in temperature-controlled rooms (21°C) on a 12-hour light/dark cycle with *ad libitum* access to food and water, nesting material and environmental enrichment consisting of tunnels and shelters. No sex differences were observed in afferent responses to acidic pH between GPR68^+/+^ and GPR68^-/-^; therefore, data from male and female mice were pooled for all subsequent analyses.

### 2.3. Reagents

Stock concentrations of Ogerin (100 mM in DMSO) and Ogerin negative control (a structurally similar inactive analogue of Ogerin; 65 mM in DMSO) were obtained from Tocris Bioscience and Sigma-Aldrich, respectively. Ogremorphin (10 mM in DMSO) was synthesized at AstraZeneca. Capsaicin (1 mM in 100% ethanol), atropine (100 mM in 100% ethanol) and nifedipine (100 mM in DMSO) were obtained from Sigma-Aldrich and bradykinin (1 mM in DMSO) was obtained from Tocris. Immediately before use, compounds were diluted to their final working concentrations in either extracellular solution (ECS) or Krebs buffer, as appropriate. 1M HCl was used to adjust the pH of ECS or Krebs buffer.

### 2.4. *In silico* analysis of human colonic biopsy bulk RNA-seq and murine colonic sensory neuron single-cell RNA-seq datasets

Bulk RNA sequencing data from colonic biopsies were obtained from a previously published dataset from Higham et al. (30) consisting of samples from patients with ulcerative colitis (UC; n = 9 biopsies from N = 9 patients), treatment-naïve Crohn’s disease (CDN; n = 8 biopsies from N = 7 patients) and treatment-refractory Crohn’s disease (CDT; n = 11 biopsies from N = 7). All IBD patients had reported abdominal pain within the four weeks prior to endoscopy.

Samples had also been collected from the colon of people with abdominal pain, but no visible inflammation, diagnosed as recurrent abdominal pain (RAP; n = 21 biopsies from N = 16) and control samples (NI; n = 14 biopsies from N = 8) obtained from individuals with no symptoms of abdominal pain or evidence of mucosal inflammation on endoscopy.

Single-cell RNA sequencing (scRNA-seq) data from murine dorsal root ganglion (DRG) neurons were acquired from Hockley et al. (31). In brief, this study performed transcriptomic profiling and unsupervised clustering of 314 retrogradely traced mouse colonic sensory neurons isolated from thoracolumbar (TL) and lumbosacral (LS) DRG (31).

### 2.5. DSS-induced acute colitis

Following baseline weight measurements on day 0, GPR68^+/+^ and GPR68^-/-^ mice of either sex received drinking water supplemented with 1.5% dextran sulphate sodium (DSS; Thermo Fisher Scientific) for 5 consecutive days, followed by normal drinking water for an additional 2 days. Control mice were provided with normal drinking water without DSS throughout the experiment. Mice were monitored daily, and disease severity was quantified using a disease activity index (DAI) as conducted previously (32). The DAI was calculated as the sum of scores for weight loss (0 = none; 1 = 1–5%; 2 = 5–10%; 3 = 10–15%; 4 = >15%), stool consistency (0 = normal; 2 = loose stool; 4 = watery diarrhoea), and presence of blood in stool (0 = none; 2 = slight bleeding; 4 = gross bleeding).

At the end of the study, mice were euthanised as described in section 2.1 and colon length and spleen weight were measured.

#### 2.5.1 Colon histology

A segment of distal colon (∼1 cm from the rectum) was dissected and fixed overnight at 4°C in 4% (w/v) formaldehyde (Sigma Aldrich). The following day, the tissue was washed in PBS and cryoprotected overnight at 4°C in 30% (w/v) sucrose (Thermo Fisher Scientific). Tissue was then embedded in Shandon M-1 Embedding Matrix (Thermo Fisher Scientific), frozen on dry ice, and stored at -80°C. Serial 20 µm sections were made using a Leica CM3000 cryostat and stored at -20°C until staining. Sections were stained with haematoxylin (0.2% w/v; Sigma Alderich; nuclei stain), and eosin (0.5% w/v; Acros Organics; cytoplasmic counterstain) as described previously (33) and mounted with glycerol. A NanoZoomer S360 and NDP.view2 (Hamamatsu) was used to image and visualise the staining, respectively.

Histopathological scoring was conducted by blinded investigators and evaluated three parameters: inflammatory cell infiltration (0 = none; 1 = increased immune nuclei in lamina propria; 2 = extension into submucosa; 3 = transmural infiltration), crypt damage (0 = intact crypts; 1–5 = increasing severity from partial crypt loss to confluent epithelial erosion), and ulceration (0 = none; 1–5 = increasing number and extent of ulcerative foci). At least two sections per animal, spaced 150 µm apart, were independently scored by two experimenters and the average score per mouse was analysed.

### 2.6. Isolation and culture of primary mouse DRG neurons

Mice were euthanised as described in section 2.1 and thoracolumbar (T12-L5) DRG were dissected, with spinal levels selected due to their innervation of the distal colon (34). Isolated DRG were transferred to 2 ml ice-cold Leibovitz L-15 medium supplemented with GlutaMAX and 2.6% (v/v) NaHCO_3_. Tissues were enzymatically digested at 37°C using type 1A collagenase (1 mg/ml, 15 min) followed by trypsin (1 mg/ml, 30 min), both prepared with 6 mg/ml bovine serum albumin (BSA). Following digestion, DRG were resuspended in 2 ml of dissociation medium (Leibovitz L-15 with GlutaMAX, 2.6% (v/v) NaHCO₃, 10% fetal bovine serum (FBS), 1.5% (v/v) glucose, and 300 U/ml penicillin / 0.3 mg/ml streptomycin). Mechanical trituration was performed with increasing force, and the resulting suspension was centrifuged at 100 g to collect the supernatant across five sequential triturations.

The pooled supernatants were centrifuged again (5 min, 100 g), pellets resuspended, and 50 µl of cell suspension was plated onto laminin-coated poly-D-lysine-coated glass-bottom dishes (MatTek, Ashland, MA). Cultures were incubated at 37°C for 2–3 hours to allow cell attachment, after which dishes were flooded with additional dissociation medium and maintained overnight at 37°C in 5% CO₂.

### 2.7. Ca^2+^ imaging

Cultured DRG neurons were incubated with 10 µM Fluo-4-AM (Invitrogen), diluted in extracellular solution (ECS) containing (in mM): 140 NaCl, 4 KCl, 1 MgCl_2_, 2 CaCl_2_, 4 D-glucose, and 10 HEPES, adjusted to pH 7.4 with NaOH and 290-310 mOsm with sucrose. Incubation was carried out for 30-45 minutes at room temperature protected from light, following which cells were gently washed with ECS and the glass-bottom dishes mounted onto the stage of an inverted Nikon Eclipse TE-2000S microscope for imaging.

Fluo-4 was excited with a 470 nm LED light source (Cairn Research, Kent, UK), and emission detected at 520 nm using Micro-manager software (v1.4; NIH). Intracellular Ca^2+^ transients were recorded using a Retiga Electro CCD camera (Photometrics, Tucson, AZ) and images were captured at 2.5 frames per second with 100 ms exposure time. A gravity-fed perfusion system (Warner Instruments, MA, USA) with 250 µm perfusion tip (Digitimer, UK) was used to continuously superfuse cells with ECS or test solutions at a rate of ∼0.5 ml/min.

#### 2.7.1 Ca^2+^ imaging protocols

To assess drug responses, cells were superfused with ECS for 10 seconds to establish baseline fluorescence, and for 4 minutes between treatments to allow for recovery. Acidic pH was applied to cells for 10-30 seconds followed by 10 seconds capsaicin (1 µM). The ECS pH was adjusted immediately preceding the experiment using HCl (1M). For antagonist studies, cells were pre-incubated for 10 minutes with 200 µl Ogremorphin (OGM; 300 nM) prior to imaging, and the inhibitor present throughout recordings. At the end of each protocol, a 10-second application of 50 mM KCl was used to depolarise neurons, confirm viability and enable fluorescence normalisation. For studies where extracellular Ca^2+^ was omitted, cells were incubated with Ca^2+^-free ECS for 10 minutes following Fluo-4 incubation. Ca^2+^ free ECS contained (in mM): 140 NaCl, 4 KCl, 2 MgCl₂, 4 glucose, 10 HEPES, and was supplemented with 1 EGTA to chelate residual Ca^2+^. To equilibrate intracellular Ca^2+^ prior to 50 mM KCl application, cells were perfused with normal ECS for 10 minutes following the application of acidic pH in Ca^2+^-free ECS.

#### 2.7.2 Ca^2+^ imaging data analysis

Image analysis was conducted using Fiji/ImageJ (NIH, MA, USA). Regions of interest (ROIs) were manually drawn around individual neurons, and mean pixel intensity (F) per frame was computed. Using custom-written scripts in RStudio, background fluorescence was subtracted, and intensity values were normalised to the maximal fluorescence elicited by 50 mM KCl (F_pos_). Neurons were considered responders if their F/F_pos_ increased by ≥0.1 during stimulation. For each condition, both the proportion of responsive neurons and the average response magnitude were compared across groups.

### 2.8. *Ex vivo* colonic afferent recording

Mice were euthanised as described in section 2.1, and the distal colon along with the associated lumbar splanchnic nerve (LSN) was isolated via laparotomy. Tissue was immediately transferred to a Sylgard-coated recording chamber maintained at 32-35°C. The colorectum was cannulated at both ends and perfused luminally (100 µl/min) and serosally (7 ml/min) with carbogenated Krebs buffer (95% O_2_, 5% CO_2_). The Krebs buffer consisted of (in mM): 124 NaCl, 4.8 KCl, 1.3 NaH_2_PO_4_, 25 NaHCO_3_, 1.2 MgSO_4_, 11.1 D-glucose, and 2.5 CaCl_2_, and was supplemented with nifedipine and atropine (both at 10 µM) to suppress smooth muscle activity.

The LSN was dissected free from surrounding connective tissue and fat, and multi-unit afferent activity was recorded using a borosilicate glass suction electrode, as conducted previously (35-37). Electrical signals were amplified (gain 5 kHz), bandpass filtered (100-1300 Hz; Neurolog, Digitimer Ltd, UK) and digitised (20 kHz; micro1401, CED, UK). Signals were visualised in real time using Spike2 software (v8.23; CED, UK), with digital filtering for 50 Hz noise (Humbug, Quest Scientific, Canada). For single-unit experiments, teased fibre recordings were performed, and individual units were identified by waveform matching to characterise single fibre properties (30, 38).

#### 2.8.1 Electrophysiology protocols

Colonic afferent preparations were left for 30-40 minutes to stabilise prior to protocol initiation. Krebs buffer was adjusted to the desired pH with HCl (1 M) immediately before application. To assess the contribution of GPR68 to colonic afferent responses to extracellular acidification, preparations from GPR68^+/+^ and GPR68^-/-^ mice were exposed to Krebs buffer at decreasing pH values (pH 6.5 to 4.0) via serosal perfusion to generate pH-response curves. A recovery period of 15-20 minutes was allowed between each stimulation to prevent desensitisation.

To further examine the role of GPR68, colonic afferent preparations were exposed to acidic Krebs buffer (pH 6.0 or 5.5) in the presence of either the GPR68 positive allosteric modulator, Ogerin (100 µM), or the selective antagonist OGM (3 µM, 30 µM). Protocols consisted of three repeated pH challenges at 20-minute intervals, with Ogerin or OGM included only during the second stimulation to assess their modulatory effects. In GPR68^-/-^ or OGM-treated groups, neuronal functionality was confirmed at the end of the protocol by applying bradykinin (1 µM), capsaicin (1 µM) and a ramp distension (0-80 mmHg).

To assess the role of GPR68 in mechanical hypersensitivity, wild-type colonic afferent preparations were exposed to a series of 5 ramp distensions at 15-minute intervals. Luminal pressure was increased to 80 mmHg (pressure transducer: Neurolog model NL108) over ∼250 seconds, which is above the threshold for visceral mechanoreceptors (39, 40). Krebs buffer at physiological or slightly acidic pH (pH 7.4 or 6.8, respectively) in the presence or absence of Ogerin (100 µM) was perfused intraluminally (Harvard Apparatus, Pump 11 Elite) between ramps 3 and 5.

#### 2.8.2 Electrophysiology data analysis

Electrophysiological activity of the LSN was analysed using Spike2 software (v8.23; CED, UK). Multi-unit spike detection was performed by applying a threshold set at twice the background noise level (30-50 µV). For protocols involving direct application of acidic Krebs buffer, spike counts were binned to calculate average firing frequency in 60-second intervals. The average firing frequency during the 6 minutes prior to acid application served as the baseline to which the peak firing rate during acid exposure was normalised. Peak changes in firing rate and area under the curve (AUC) were calculated for each pH challenge and used to compare responses across genotypes and treatment conditions. For Ogerin and OGM datasets, the second response (in the presence of the modulators) was compared to the first response (pre-modulator incubation).

For ramp distension protocols, changes in firing rate were quantified relative to baseline firing (3-minutes prior to distension). Nerve activity was assessed at 5 mmHg increments, and from these data, both the peak firing rate (at 80 mmHg) and AUC were calculated. The response to ramp 5 (post-drug) is expressed as a percentage of the response to ramp 3 (pre-drug). A post-treatment response of ∼100% indicates no change in distension-evoked firing.

### 2.9. Statistics

Normality was assessed and corresponding parametric or non-parametric statistical tests were conducted. Data are presented as mean ± standard error of the mean (SEM) and statistical significance was defined as P ≤ 0.05. For Ca^2+^ imaging experiments, ‘n’ refers to the number of individual culture dishes analysed, while ‘N’ denotes the total number of animals from which these cultures were derived. In electrophysiological recordings, N represents the number of animals used. All statistical analyses were conducted using GraphPad Prism (version 10.4.2; GraphPad Software, San Diego, CA, USA). Schematics are included throughout the manuscript to aid interpretation of experimental design (created with BioRender.com and ChemDraw (v21.1.2)).

## 3. Results

### 3.1 *In silico* and *in vivo* analyses implicate GPR68 in IBD pathology and reveal co-expression with *Trpv1* in murine colon-projecting sensory neurons

*In silico* analysis of transcript expression in colonic biopsies revealed a 3-fold increase in *GPR68* mRNA in samples from individuals with ulcerative colitis (UC) compared to noninflamed controls (NI) (P = 2.8 x 10^-4^, one-way ANOVA with Benjamini-Hochberg post-hoc test; **Fig. 1A**). *GPR68* expression was also elevated in both drug-naïve and drug-treated Crohn’s disease samples (CDN and CDT, respectively) (NI vs CDN: P = 2.6 x 10^-4^; NI vs CDT: 5.8 x 10^-3^; one-way ANOVA with Benjamini-Hochberg post-hoc test; **Fig. 1A**; data redrawn from Higham et al. (30)).

**Figure 1.**
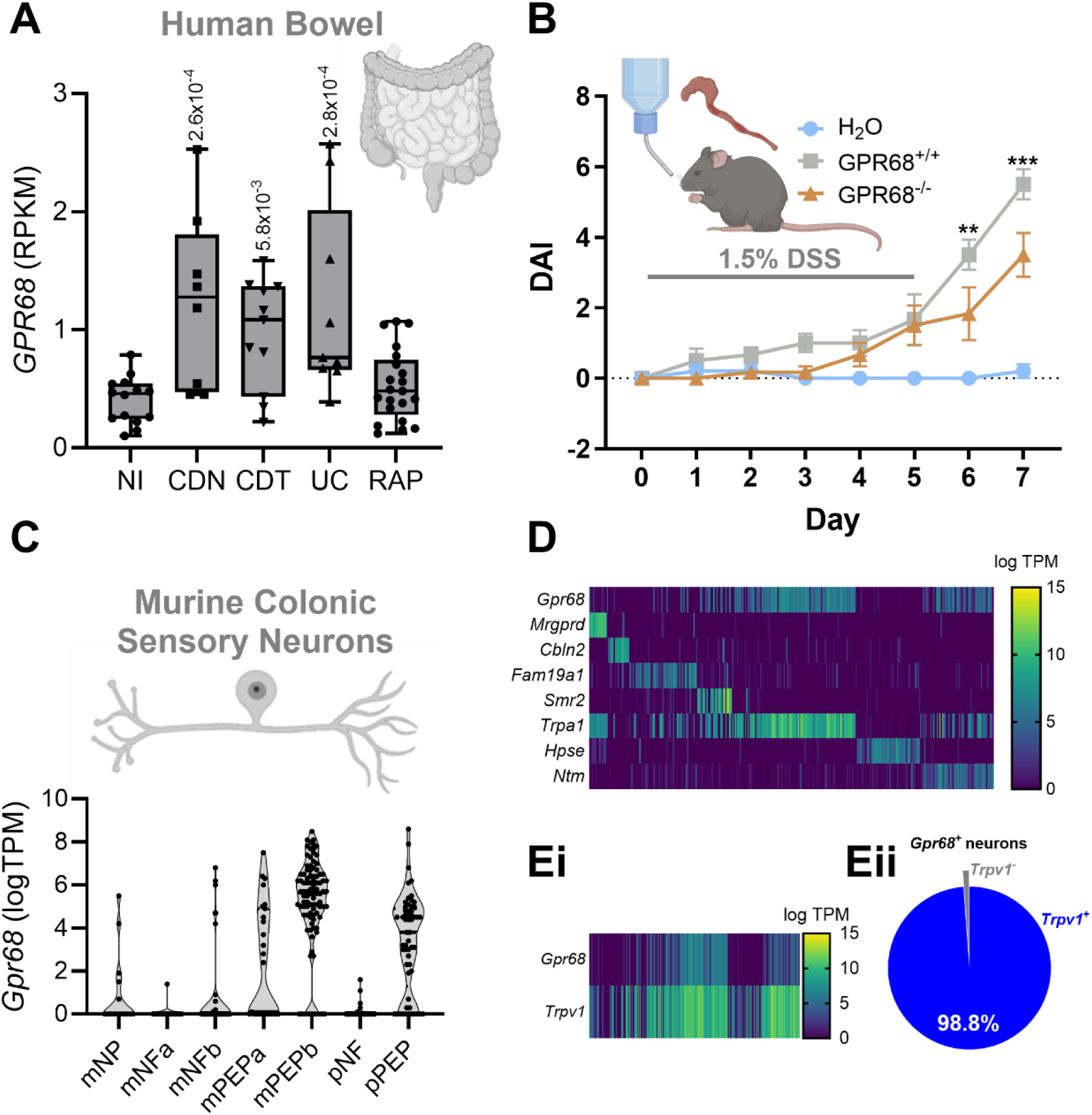
GPR68 is upregulated in bowel tissue from individuals with colitis and enriched in murine colon-innervating sensory neurons. (A) Expression (RPKM, Reads Per Kilobase Million) of *GPR68* in colonic biopsies taken from people with no inflammation (NI), ulcerative colitis (UC), drug-naïve and drug-treated Crohn’s disease (CDN and CDT, respectively) and recurrent abdominal pain (RAP). Median, interquartile range (box), and full range (whiskers) is shown. (*GPR68* CDN vs NI, P = 2.6 x 10^-4^; CDT vs NI, P = 5.8 x 10^-3^; UC vs NI, P = 2.8 x 10^-4^; one-way ANOVA (main effect: P = 6.53 x 10^-5^; F(4, 58) = 3.579) with Benjamini-Hochberg post-hoc test). Data redrawn from Higham et al. (30). (B) Disease activity index (DAI) over the course of 1.5%-induced colitis in (GPR68^+/+^ vs GPR68^-/-^ Day 6, P = 0.003**; Day 7, P = 0.0003***; two-way ANOVA (main effect: P = <0.0001; F(2, 112) = 40.57) with Tukey’s post-hoc test). (C) Violin plot showing the expression (logTPM) of *Gpr68* in the murine colonic afferent populations identified by Hockley et al. (31). (D) Heatmap showing the expression (logTPM) of *Gpr68* and markers of each of the murine colonic afferent populations. Heatmap (Ei) and Chart (Eii) showing the fraction of *Gpr68*-expressing sensory neurons which co-express *Trpv1*. Data redrawn from Hockley et al. (31).

To investigate the functional role of GPR68 in colitis, we used a 1.5% DSS-induced colitis model (32). GPR68^-/-^ mice exhibited significantly reduced disease activity compared to wild-type controls on days 6 and 7 (GPR68^+/+^ vs GPR68^-/-^ Day 6, P = 0.003; Day 7, P = 0.0003; two-way ANOVA with Tukey’s post-hoc test; N = 6; **Fig. 1B**). No significant differences were observed in colon length, spleen weight or histological inflammation scores (**Supplemental Fig. 2**).

Further supporting a role for GPR68 in colitis-associated nociception, we found that *Gpr68* expression was enriched in peptidergic subpopulations of colonic sensory neurons (**Figs. 1C-D**) and showed high co-expression with *Trpv1*, a marker of nociceptive afferents (**Fig. 1E**). Furthermore, comparison of GPR68 with other proton-sensing GPCRs (GPR4, GPR65 and GPR132) revealed that although expression of each receptor was upregulated to varying degrees in colitic tissue, GPR68 was the most highly expressed proton-sensing GPCR in colonic sensory neurons, highlighting its potential contribution to both inflammation and visceral pain (**Supplemental Fig. 1**).

### 3.2 GPR68 contributes to acid-evoked intracellular Ca^2+^ responses in DRG sensory neurons

Given the expression of *Gpr68* in mouse colonic sensory neurons, we next sought to investigate the contribution of GPR68 to acid-induced intracellular Ca^2+^ mobilisation ([Ca^2+^]_i_) in cultured thoracolumbar DRG neurons. We first characterised responses to acidic pH and capsaicin in neurons isolated from GPR68^+/+^ and GPR68^-/-^ mice under normal extracellular Ca^2+^ conditions observing no significant difference in either the peak response or the proportion of DRG neurons responding across the pH range (pH 6.5-5.0) (**Figs. 2A-E**) and to capsaicin (**Figs. 2I-L**). In both genotypes, acid-sensitive neurons were significantly smaller than non-responders (P = <0.0001, Mann Whitney tests; **Figs. 2F-G**) and exhibited high capsaicin co-sensitivity (pH 6.0: 84.5 ± 4.9% and 88.8 ± 3.0% for GPR68^+/+^ and GPR68^-/-^, respectively), consistent with the activation of *Trpv1*-positive sensory neurons (**Figs. 2A-E).**

**Figure 2.**
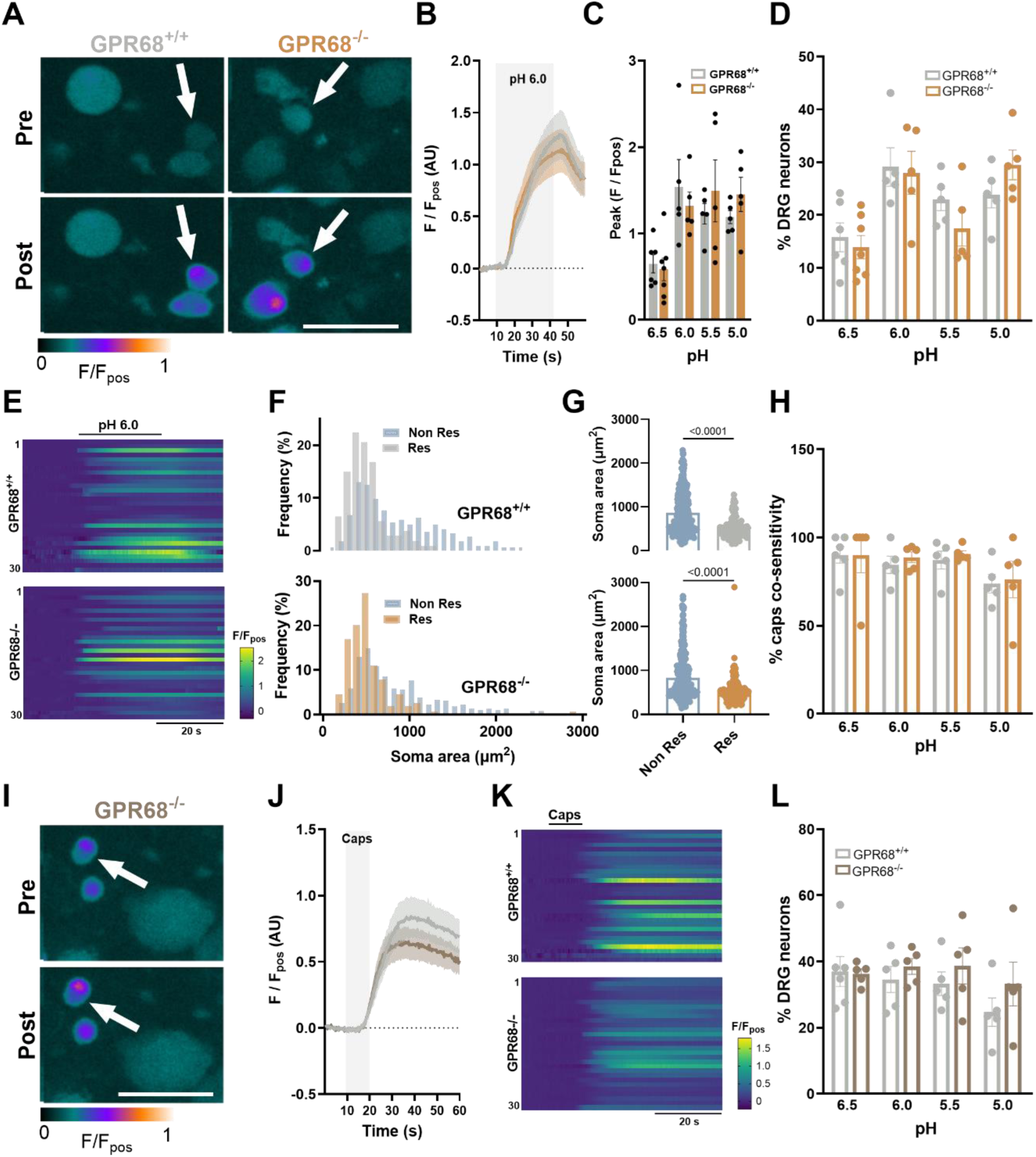
Characterisation of acid- and capsaicin-evoked [Ca^2+^]_i_ mobilisation in DRG neurons isolated from GPR68^+/+^ and GPR68^-/-^ mice. (A) False-coloured images showing Fluo-4 fluorescence during the application of normal pH 6.0 ECS to DRG neurons isolated from GPR68^+/+^ and GPR68^-/-^ mice. White arrows highlight cells responding to pH 6.0. Scale bar: 50 µM. (B) Averaged traces of pH 6.0 response and (C) peak responses across the pH range in DRG neurons isolated from GPR68^+/+^ and GPR68^-/-^ mice (GPR68^+/+^ vs GPR68^-/-^ P = 0.9984, 0.9117, 0.8385, 0.8443 (left to right); one-way ANOVA (main effect: P = 0.0031, F(7, 35) = 3.901) with Sidak’s multiple comparisons test; n = 5-7 dishes from N = 5 animals). (D) The proportion of GPR68^+/+^ and GPR68^-/-^ DRG neurons responding to pH 6.5-5.0. (GPR68^+/+^ vs GPR68^-/-^ P = 0.9808, 0.9984, 0.6238, 0.5866 (left to right); one-way ANOVA (main effect: P = 0.0008, F(7, 35) = 4.701) with Sidak’s multiple comparisons test; n = 5-7 dishes from N = 5 animals). (E) Heatmap showing Fluo-4 fluorescence during the application of pH 6.0 for 30 randomly selected DRG neurons for 30 seconds isolated from GPR68^+/+^ (top) and GPR68^-/-^ (bottom) mice. (F) Cell size frequency distribution and (G) scatter dot plot of average soma areas (µm^2^) of responders and non-responders to pH 6.0 in both genotypes (GPR68^+/+^ P = <0.0001 (U = 10075); GPR68^-/-^ P = <0.0001 (U = 17675); Mann-Whitney Test; n = 445 and n = 548 cells, respectively). (H) The proportion of pH 6.0-responding DRG neurons also co-sensitive to capsaicin (GPR68^+/+^ vs GPR68^-/-^ P = >0.9999, 0.9823, 0.9923, 0.9983 (left to right); one-way ANOVA (main effect: P = 0.3866, F(7, 33) = 1.099) with Sidak’s multiple comparisons test; N = 5-6). (I) False-coloured images showing Fluo-4 fluorescence in GPR68^-/-^ DRG neurons in response to capsaicin (1 µM). (J) Averaged traces and heatmaps of 30 selected GPR68^+/+^ and GPR68^-/-^ DRG neurons (K) portraying responses to capsaicin. (L) The proportion of GPR68^+/+^ and GPR68^-/-^ DRG neurons responding to capsaicin (GPR68^+/+^ vs GPR68^-/-^ P = 0.9520, 0.8642, 0.5481, >0.9999 (left to right); one-way ANOVA (main effect: P = 0.4100, F(7, 33) = 1.061) with Sidak’s multiple comparisons test; n = 5-6 dishes from N = 5 animals).

While no differences in pH responses were observed under normal extracellular Ca^2+^ conditions, repeating experiments in Ca^2+^ free buffer, thereby restricting [Ca^2+^]_i_ mobilisation to intracellular stores (confirmed by loss of response to capsaicin; **Supplemental Figure 3**), revealed a significant decrease in the proportion of DRG neurons responding to pH 6.5 and pH 6.0 in neurons isolated from GPR68^-/-^ mice compared with wild-type mice or following pre-treatment with the GPR68 antagonist, Ogremorphin in wild-type mice (OGM; pH 6.5: Control vs OGM: P = 0.0051; Control vs GPR68^-/-^: P = 0.0274; one-way ANOVA with Dunnett’s multiple comparisons test; N = 5-8; **Figs. 3A-F**). Additionally, the magnitude of the response in the remaining acid-sensitive neurons was also significantly reduced in cells from GPR68^-/-^ mice or OGM-treated cells (pH 6.5: Control vs OGM: P = 0.0499; Control vs GPR68^-/-^: P = 0.0340; one-way ANOVA with Dunnett’s multiple comparisons test; N = 5-8; **Figs. 3A-F**). These results confirm that GPR68 contributes to acid-induced [Ca^2+^]_i_ mobilisation in DRG sensory neurons. Experiments were performed using pH 6.5 and pH 6.0 as these concentrations of protons have previously been reported to elicit robust GPR68 activity (17) with minimal TRPV1 activity (41).

**Figure 3.**
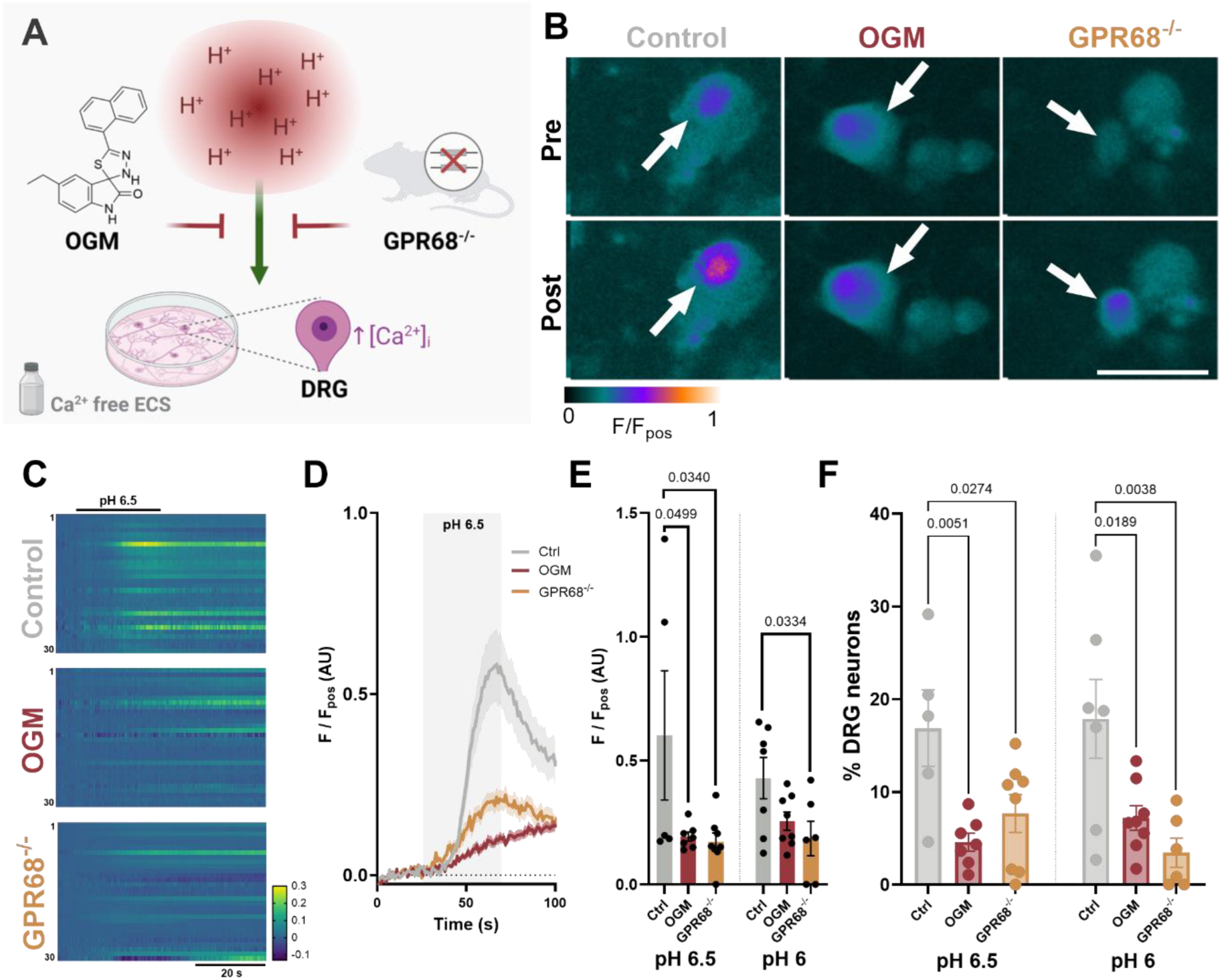
GPR68 contributes to acid-evoked [Ca^2+^]_i_ increase in DRG sensory neurons following removal of external Ca^2+^. (A) Schematic of the experimental paradigm assessing DRG neuron responses to acidic pH in Ca^2+^-free ECS. (B) False-coloured images and (C) Heatmaps (30 randomly selected neurons) showing Fluo-4 fluorescence during the application of Ca^2+^-free ECS adjusted to pH 6.5 in GPR68^+/+^ and GPR68^-/-^ DRG neurons and following pre-treatment with GPR68 antagonist OGM. White arrows highlight cells responding to pH 6.5. Scale bar: 50 µM. (D) Averaged response profiles and peak responses (E) to acidic pH across treatment conditions (pH 6.5 Control vs OGM: P = 0.0499; Control vs GPR68^-/-^: P = 0.0340; one-way ANOVA (main effect: P = 0.0403; F(2, 17) = 3.903) with Dunnett’s multiple comparisons test. pH 6.0 Control vs OGM: P = 0.1101; Control vs GPR68^-/-^: P = 0.0334; one-way ANOVA (main effect: P = 0.0451; F(2, 18) = 3.7) with Dunnett’s multiple comparisons test; n = 5-8 dishes from N = 5-6 animals) (F) The proportion of GPR68^+/+^ and GPR68^-/-^ DRG neurons responding to pH 6.5 and pH 6.0 and following pre-treatment with OGM (pH 6.5 Control vs OGM: P = 0.0051; Control vs GPR68^-/-^: P = 0.0274; one-way ANOVA (main effect: P = 0.0085; F(2, 17) = 6.392) with Dunnett’s multiple comparisons test. pH 6.0 Control vs OGM: P = 0.0189; Control vs GPR68^-/-^: P = 0.0038; one-way ANOVA (main effect: P = 0.0048; F(2, 18) = 7.281) with Dunnett’s multiple comparisons test; n = 5-8 dishes from N = 5-6 animals).

### 3.3 Colonic afferent responses to acidic pH are attenuated in GPR68^-/-^ mice and inhibited in wild-type mice by GPR68 inhibition

Having established the presence of GPR68 mediated acid-evoked stimulation of DRG sensory neurons, we next investigated the contribution of GPR68 to acid-evoked stimulation of colonic afferents using the LSN preparation.

Repeat application of acidic Krebs buffer (pH 6.5-4.0) elicited a robust increase in colonic afferent activity in tissue from GPR68^+/+^ mice which was absent in tissue from GPR68^-/-^ mice (P = <0.0001; two-way ANOVA; N = 9-11; **Figs. 4A-B**). This reduction was reflected in both the peak change in nerve activity and area under the curve (AUC) of the colonic afferent response (peak activity: pH 6.0: 1.277 ± 0.339 spikes/s, P = 0.0074; pH 4.0: 4.452 ± 1.82 spikes/s, P = 0.0008; AUC: pH 5.5 P = 0.003; Mann-Whitney tests; N = 9-11; **Figs. 4C-D**).

**Figure 4.**
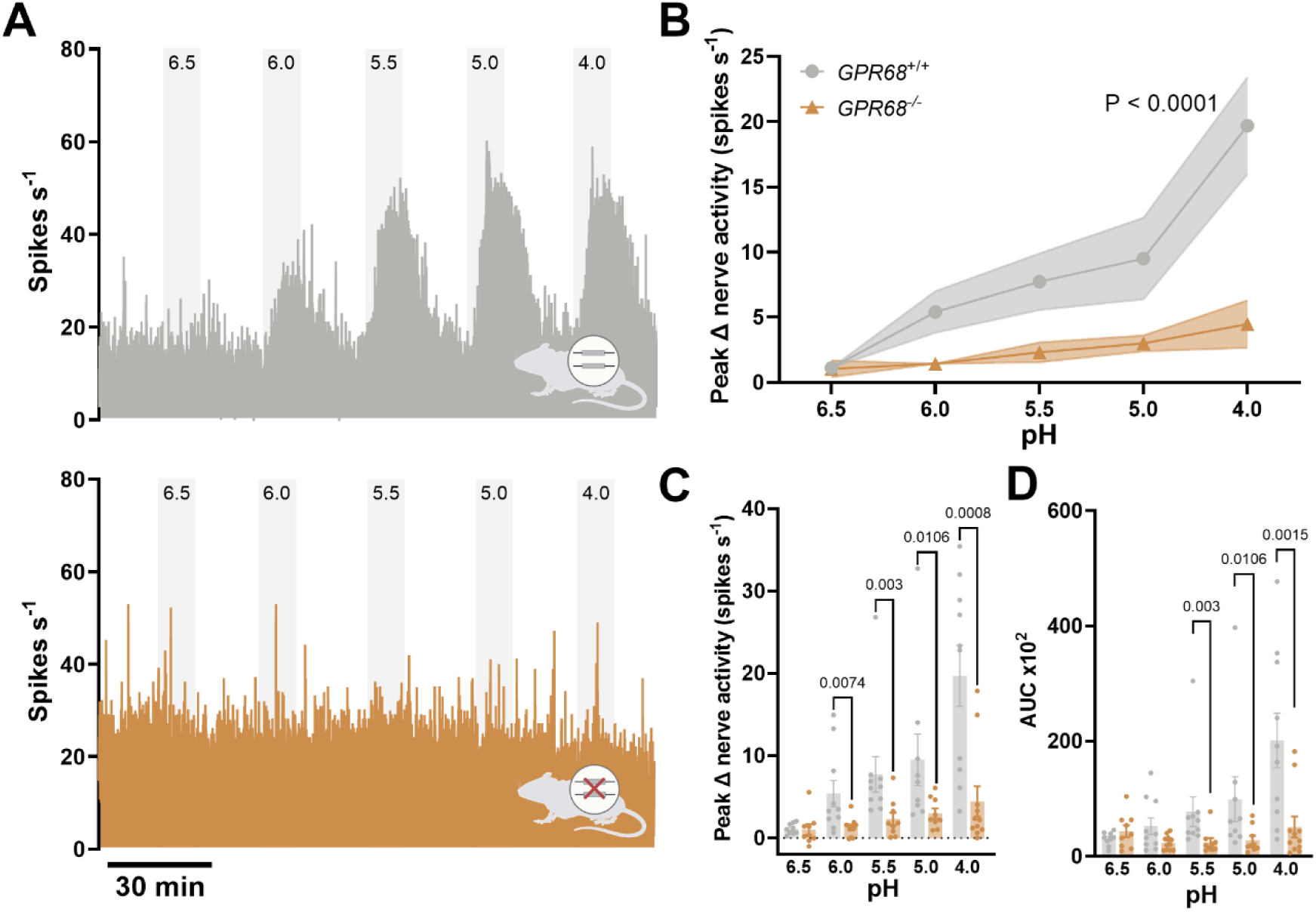
Colonic afferents from GPR68^-/-^ mice exhibit reduced acid-sensitivity. (A) Representative rate histograms and (B) mean response profiles of GPR68^+/+^ (grey) and GPR68^-/-^ (orange) colonic afferent activity in response to pH 6.5-4.0; shaded area is SEM (GPR68^+/+^ vs GPR68^-/-^ main effect: P = <0.0001; two-way ANOVA; F(1, 87) = 26.47). (C) Peak afferent activity at each pH (GPR68^+/+^ vs GPR68^-/-^ unpaired t-test (or non-parametric equivalent); pH 6.5, P = 0.9126, t, df = 0.1115, 16; pH 6.0, P = 0.0074 (U = 18); pH 5.5, P = 0.003 (U = 10); pH 5.0, P = 0.0106 (U = 12); pH 4.0, P = 0.0008 (U = 10); N = 9-11 animals). (D) Area under the curve (AUC) for afferent responses across the pH range (GPR68^+/+^ vs GPR68^-/-^ unpaired t-test (or non-parametric equivalent); pH 6.5, P = 0.2430, t, df = 1.212, 16; pH 6.0, P = 0.1145 (U = 32); pH 5.5, P = 0.003 (U = 10); pH 5.0, P = 0.0106 (U = 12); pH 4.0, P = 0.0015 (U = 12); N = 9-11 animals).

Importantly no significant differences were seen in the LSN response to ramp distension (P = 0.7861; unpaired t test; N = 7-8), bradykinin (P = 0.1785 unpaired t test; N = 8-10) or capsaicin (P = 0.5116; Mann Whitney test; N = 10-11) between GPR68^+/+^ and GPR68^-/-^ preparations, confirming that the loss of acid-evoked colonic afferent activity was stimulus-specific and indicating that responses to noxious stimuli and GPCR signalling in general remained intact (**Fig. 5**). Furthermore, acid-evoked responses in tissue from GPR68^+/-^ mice were comparable to those in wild-type controls (P = 0.5185; two-way ANOVA; N = 9-10; **Supplemental Figure 4**).

**Figure 5.**
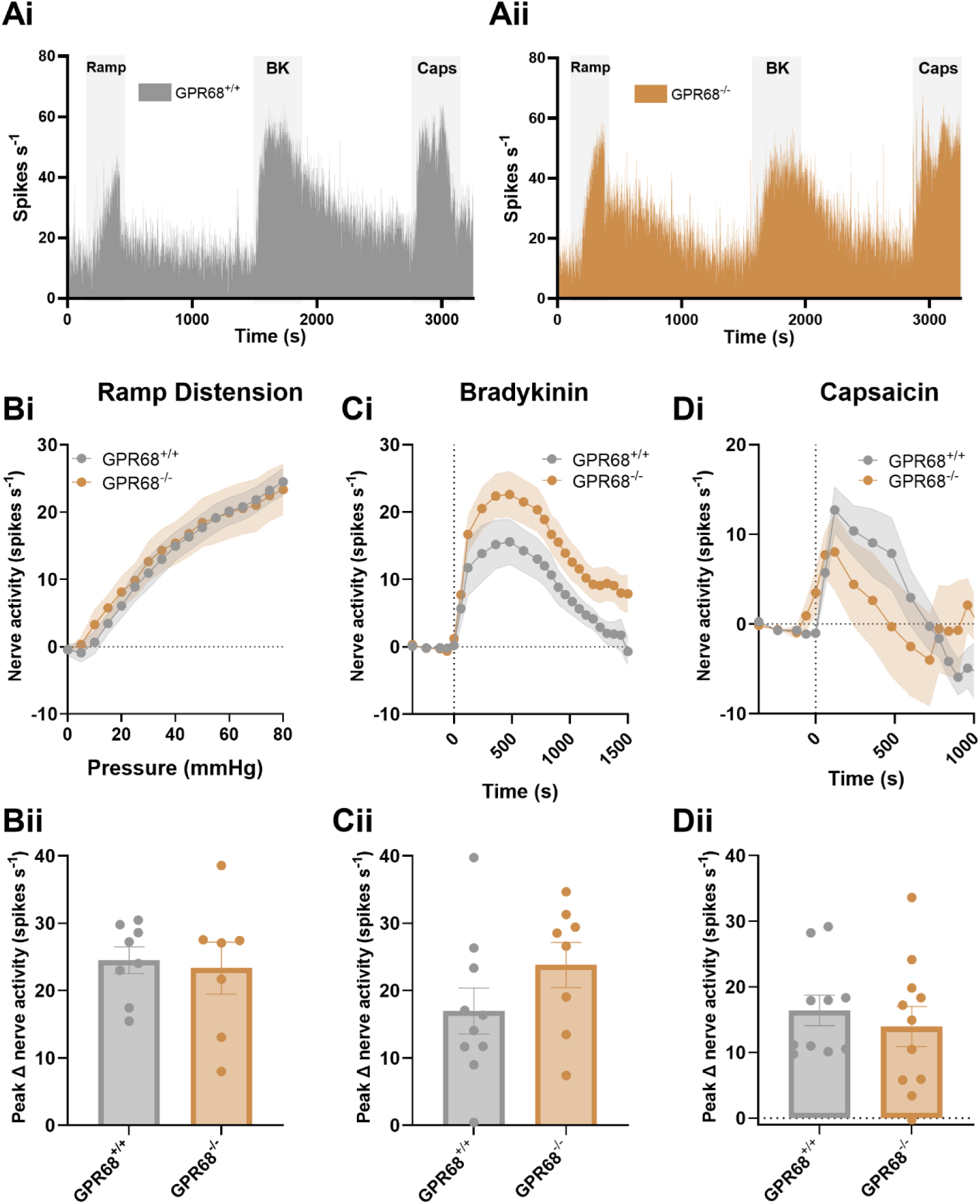
GPR68^+/+^ and GPR68^-/-^ colonic afferent activity is comparable in response to mechanical and chemical (bradykinin and capsaicin) stimuli. (A) Representative rate histogram of (i) GPR68^+/+^ and (ii) GPR68^-/-^ colonic afferent responses to ramp distension (Ramp), bradykinin (BK; 1 µM) and capsaicin (Caps; 1 µM). (Bi) Response profile (P = 0.8246, two-way ANOVA; F(1, 13) = 0.05113; N = 7-8) and (Bii) peak change in afferent activity at 80 mmHg in response to ramp distension (P = 0.7861 (t, df = 0.2770, 13), unpaired t test; N = 7-8). (Ci) Response profile (P = 0.0947, two-way ANOVA; F(1, 15) = 3.182; N = 8-10) and (Cii) peak change in afferent activity in response to bradykinin stimulation (P = 0.1785 (t, df = 1.407, 16), unpaired t test; N = 8-10). (Di) Response profile (P = 0.7187, two-way ANOVA; F(1, 19) = 0.1337; N = 10-11) and (Dii) peak change in nerve activity in response to capsaicin (1 µM) stimulation (P = 0.5116, Mann-Whitney test (U = 45); N = 10-11).

Complementary to the data obtained using transgenic knockout mice, pharmacological blockade of GPR68 with the selective GPR68 antagonist, OGM (**Fig. 6A-B**) significantly attenuated the peak afferent response (vehicle vs 3 µM OGM, P = 0.0183; vehicle vs 30 µM OGM, P = 0.0044; one-way ANOVA with Dunnett’s multiple comparisons test; N = 6; **Fig. 6C**) and area under the curve for afferent activity compared to vehicle pre-treatment following application of pH 5.5 acidified buffer (vehicle vs 3 µM OGM, P = 0.0081; vehicle vs 30 µM OGM, P = 0.0057; one-way ANOVA with Dunnett’s multiple comparisons test; N = 6; **Fig. 6D**).

**Figure 6.**
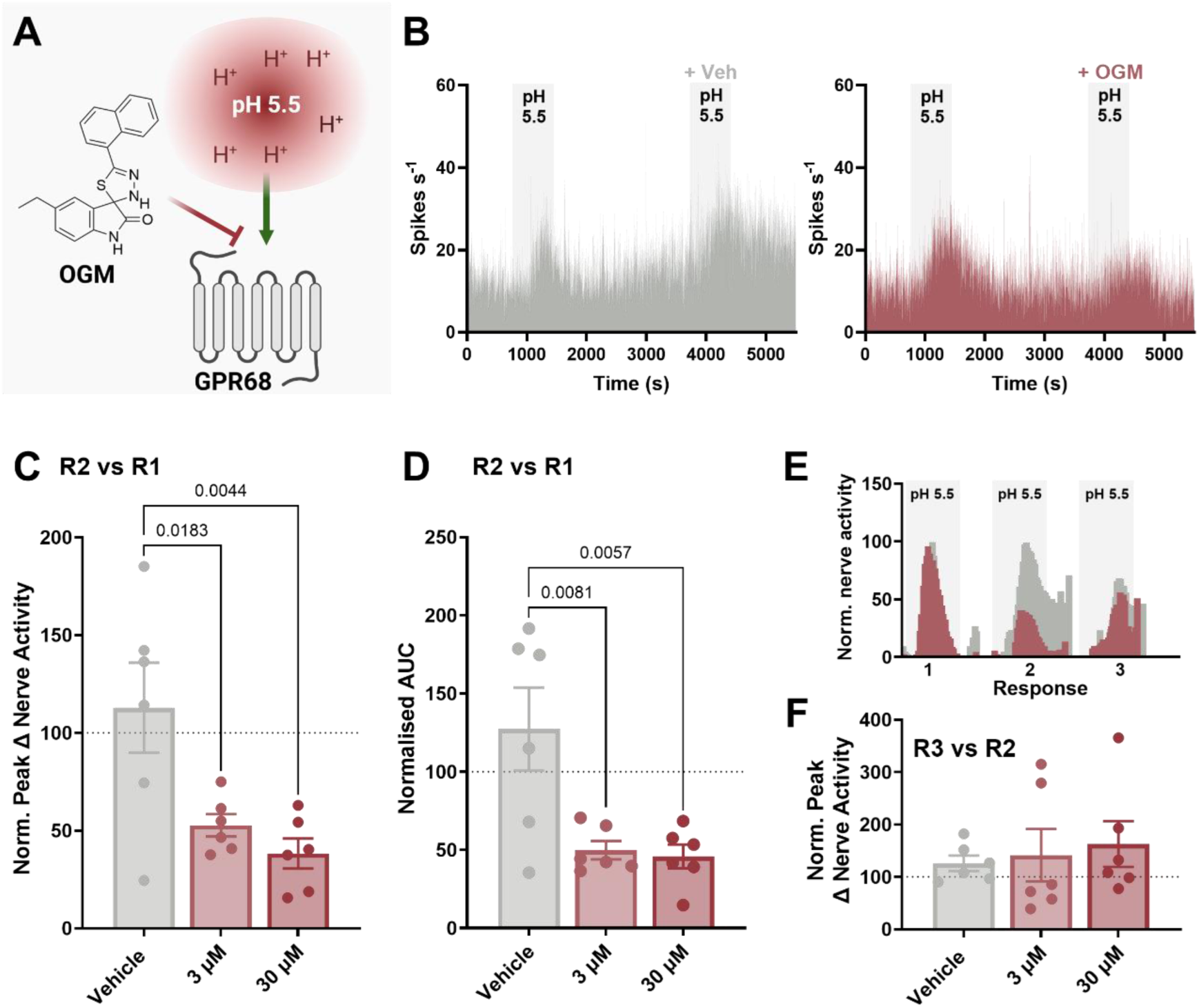
GPR68 antagonist, OGM, attenuates colonic afferent responses to acid. (A) Schematic of the experimental paradigm assessing colonic afferent responses to pH 5.5 in the presence and absence of the GPR68 antagonist, OGM. (B) Representative rate histograms of repeated stimulation with pH 5.5. The 2^nd^ response consisted of pretreatment with vehicle control (DMSO) (left) or OGM (right). (C) Peak change in afferent activity and (D) AUC of response 2 (in the presence of OGM or vehicle) expressed as a percentage of the peak response 1 (pH 5.5 alone) (Peak change in afferent activity: vehicle vs 3 µM OGM, P = 0.0183; vehicle vs 30 µM OGM, P = 0.0044; one-way ANOVA (main effect: P = 0.0054; F(2, 15) = 7.537) with Dunnett’s multiple comparison test) (AUC: vehicle vs 3 µM OGM, P = 0.0081; vehicle vs 30 µM OGM, P = 0.0057; one-way ANOVA (main effect: P = 0.0045; F(2, 15) = 7.928) with Dunnett’s multiple comparison test). (E) Averaged response profiles to pH 5.5 before (response 1), during (response 2), and after (response 3) treatment with vehicle (grey) or 3 µM OGM (red). (F) Peak change in afferent activity of response 3 normalised to peak response 2 (pH 5.5 with OGM or vehicle) (vehicle vs 3 µM OGM, P = 0.7739; vehicle vs 30 µM OGM, P = >0.9999; Kruskal-Wallis test (main effect: P = 0.4657; H = 1.626) with Dunnett’s multiple comparison test) (N = 6 animals per group).

Notably, repeat applications of pH 5.5 evoked consistent responses in control LSN recordings, with no significant difference observed across the three applications, confirming the reproducibility of the stimulus (comparison not shown). Furthermore, a recovery of the pH response during the third stimulation was not observed in OGM-treated tissue (vehicle vs 3 µM OGM, P = 0.7739; vehicle vs 30 µM OGM, P = >0.9999; Kruskal-Wallis test with Dunnett’s multiple comparison test; N = 6; **Figs. 6E-F**), suggesting sustained inhibition by OGM.

Comparable to studies performed in tissue from GPR68^-/-^ mice, responses to ramp distension (P = 0.6056; N = 3-6), bradykinin (P = 0.4315; N = 5-6) and capsaicin (P = 0.1959; N = 6) were similar between treatment groups (one-way ANOVA main effect; **Supplemental Figure 5**).

### 3.4 Positive allosteric modulation of GPR68 potentiates colonic afferent responses to acid and noxious mechanical distension

To further explore the functional consequences of GPR68 activation, we next examined the effects of Ogerin, a selective positive allosteric modulator of GPR68 (42), on colonic afferent responses to acidic and mechanical stimuli.

Consistent with a role for GPR68 in acid evoked colonic afferent activation, pre-treatment with Ogerin, but not the Ogerin negative control (ONC), significantly increased the peak change and AUC of colonic afferent responses to pH 6.0 Krebs (**Figs. 7A-B**) compared to vehicle control, this effect was no longer observed in colonic afferent recordings made from GPR68^-/-^ mice confirming the specificity of Ogerin for GPR68 receptor activity (Peak: Ogerin vs ONC, P = 0.0005; Ogerin vs Vehicle, P = 0.0019; Ogerin vs GPR68^-/-^, P = 0.0002; one-way ANOVA with Dunnett’s multiple comparisons test; AUC: Ogerin vs ONC, P = 0.0047; Ogerin vs Vehicle, P = 0.0014; Ogerin vs GPR68^-/-^, P = 0.0237; one-way ANOVA with Dunnett’s multiple comparisons test; N = 5-6; **Fig. 7C-D**). Furthermore, single-unit LSN recordings showed that Ogerin potentiated the response to acidic pH in a mechanosensitive fibre responsive to colonic distension at pressures >20 mmHg (**Fig. 7E**).

**Figure 7.**
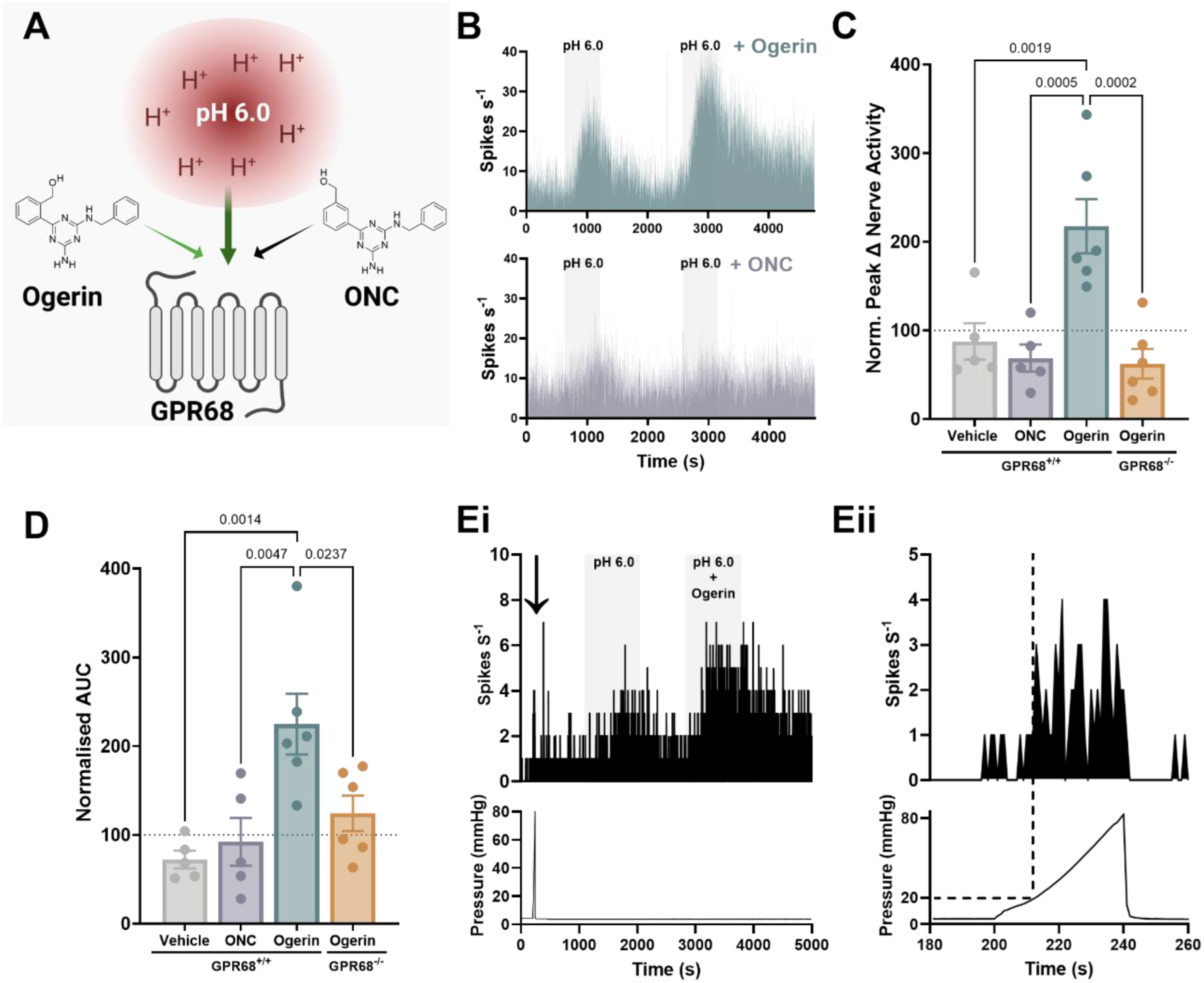
The GPR68 PAM Ogerin potentiates colonic afferent responses to acid. (A) Schematic of the experimental paradigm assessing colonic afferent responses to pH 6.0 following treatment with the GPR68 positive allosteric modulator (PAM), Ogerin, or its inactive analogue ONC (Ogerin negative control). (B) Representative rate histograms of repeated afferent stimulation with pH 6.0. The 2^nd^ response consisted of pretreatment with either Ogerin (top) or ONC (bottom). (C) Peak change in afferent activity of response 2 (post-drug application) normalised to response 1 (pre-drug application) for vehicle, ONC or Ogerin (in GPR68^+/+^ and GPR68^-/-^ mice) treated groups (Ogerin vs ONC, P = 0.0005; Ogerin vs Vehicle, P = 0.0019; Ogerin vs GPR68^-/-^, P = 0.0002; one-way ANOVA (main effect: P = 0.0002; F(3, 18) = 11.36); with Dunnett’s multiple comparisons test; N = 5-6 animals per group). (D) AUC of response 2 (post-drug application) normalised to response 1 (pre-drug application) for vehicle, ONC or Ogerin (in GPR68^+/+^ and GPR68^-/-^ mice) treated groups (Ogerin vs ONC, P = 0.0047; Ogerin vs Vehicle, P = 0.0014; Ogerin vs GPR68^-/-^, P = 0.0237; one-way ANOVA (main effect: P = 0.002; F(3, 18) = 7.367); with Dunnett’s multiple comparisons test; N = 5-6 animals per group) (Ei) Representative rate histogram (top) and corresponding pressure trace (bottom) from a single-unit recording showing afferent responses to ramp distension (0-80 mmHg) (indicated by arrow), followed by perfusion with pH 6 alone then pH 6 with Ogerin. (Eii) Expanded view of the response to luminal distension indicated with arrow in (Ei); dotted line shows increased afferent firing at luminal pressures exceeding 20 mmHg (N = 1).

To further elucidate the contribution of GPR68 to the sensitisation of mechanosensitive colonic afferents we investigated the effect of Ogerin on colonic afferent responses to luminal distension with pH 7.4 Krebs buffer and weakly acidified pH 6.8 Krebs buffer (**Figs. 8A-B**). Consistent with its role as a positive allosteric modulator of GPR68, pre-treatment with Ogerin (100 µM) had no significant effect on the colonic afferent firing response to ramp distension with pH 7.4 buffer (pH 7.4 vs pH 7.4 with Ogerin, P = 0.8263; one-way ANOVA with Dunnett’s multiple comparisons test; N = 5-6; **Figs. 8C-D**). However, Ogerin significantly enhanced the afferent response to colonic distension with pH 6.8 buffer at noxious distending pressures (75 mmHg P = 0.0236; 80 mmHg P = 0.0314; repeated measures ANOVA with Dunnet’s multiple comparisons test; N = 5-6; **Fig. 8C**), such that a significant increase in peak afferent firing during ramp distension was observed (pH 7.4 vs pH 6.8 with Ogerin; P = 0.0330; N = 5-6; **Fig. 8D**). Luminal distension with pH 6.8 buffer alone produced a comparable increase in colonic afferent activity to distension with pH 7.4 buffer.

**Figure 8.**
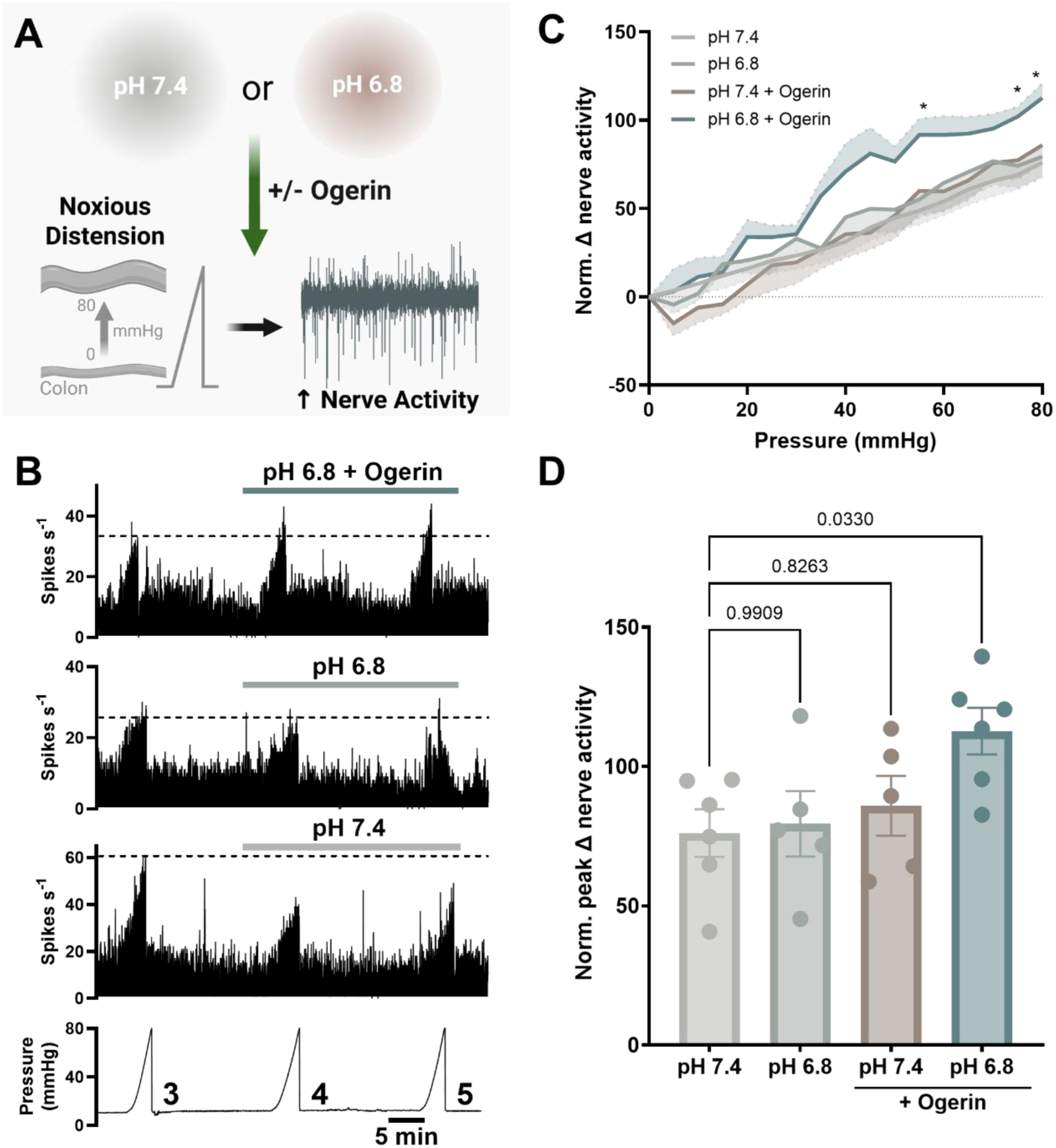
The GPR68 PAM Ogerin potentiates colonic afferent responses to noxious mechanical distension. (A) Schematic of experimental paradigm assessing the effects of the GPR68 PAM, Ogerin, on colonic afferent responses to ramp distension. (B) Representative rate histograms and corresponding pressure traces from recordings showing afferent responses to ramp distensions 3-5 in pH 7.4, 6.8 and pH 6.8 + Ogerin (100 µM) groups. The dotted line highlights the sensitisation of afferent responses to distension in the pH 6.8 + Ogerin group. (C) Afferent response profiles of ramp distension 5 (0-80 mmHg) in pH 7.4 and pH 6.8 groups with and without Ogerin, normalised (norm.) to ramp distension 3 (*P < 0.05; two-way repeated measures ANOVA (main effect: P = 0.0602; F(3, 18) = 2.957) with Dunnett’s multiple comparisons test). Shaded area is SEM. (D) Peak change in nerve activity at 80 mmHg during Ramp 5 (post-drug application), expressed as a percentage of Ramp 3 (pre-drug application), in pH 7.4 and pH 6.8 groups with and without Ogerin (pH 7.4 vs pH 6.8 with Ogerin, P = 0.0330; one-way ANOVA (main effect: P = 0.0521; F(3, 18) = 3.113) with Dunnett’s multiple comparisons test) (N = 5-6 animals).

## 4. Discussion

Abdominal pain is a leading cause of morbidity in individuals with IBD, often persisting during remission, underscoring the limitation of current therapies for the resolution of pain (43). A critical unmet clinical need therefore exists for the development of novel treatment strategies which not only target inflammation but also nociceptive signaling. To address this, we investigated the contribution of GPR68, a proton-sensing GPCR implicated in intestinal inflammation and fibrosis (21, 23, 25), to the activation of colonic afferents and DRG sensory neurons.

Consistent with reports of a pro-inflammatory role for GPR68 in colitis (22, 25), we showed that GPR68^-/-^ mice exhibited significantly less disease activity following DSS-induced acute colitis, and *in silico* analysis of our reported RNA-sequencing data revealed increased expression of GPR68 alongside other proton-sensing GPCRs in colonic biopsies from individuals with IBD (30). These data support the potential utility of GPR68 antagonists for the treatment of inflammation in IBD. Furthermore, transcriptomic analysis of murine colon-projecting sensory neurons showed that GPR68 expression was markedly higher than other proton-sensing GPCRs and was particularly enriched in *Trpv1*-expressing peptidergic nociceptors (31). These observations highlight a unique opportunity to target GPR68 to suppress both nociceptor signaling and inflammation in colitis.

To better understand the contribution of GPR68 to sensory signaling, we examined DRG neuron responses to acidic pH in Ca^2+^-free conditions, thereby isolating intracellular Ca^2+^ release. We observed a significant reduction in the amplitude and proportion of DRG neurons responding to weakly acidic pH (pH 6.0–6.5) in preparations from GPR68^-/-^ mice or following pre-treatment of DRG neurons with the GPR68 antagonist, Ogremorphin. Prior studies similarly demonstrate that acid-induced intracellular Ca^2+^ fluxes are diminished in cells expressing mutant GPR68 (44), and that knockdown of GPR68 in airway smooth muscle cells attenuates acid-evoked Ca^2+^ mobilization (45, 46). Our findings are consistent with the known G_q_-coupling of GPR68 signaling to acid (17), which liberates Ca^2+^ from intracellular stores via IP_3_ receptors following G_q_ mediated phospholipase C activation and formation of IP_3_ from phosphatidylinositol 4, 5-bisphosphate (PIP_2_). The marked reduction in acid-evoked responses in DRG neurons isolated from GPR68^-/-^ mice or following pre-treatment with Ogremorphin, is also in line with GPR68 being the most highly expressed proton-sensing GPCR in colon-projecting sensory neurons (31). The presence of a small residual population of DRG neurons exhibiting intracellular Ca^2+^ responses to acid in Ca^2+^-free conditions further suggests the involvement of other proton sensitive pathways capable of mobilizing intracellular Ca^2+^ in sensory neurons.

We observed comparable responses to capsaicin and no difference in acid-evoked responses in DRG neurons isolated from GPR68^+/+^ and GPR68^-/-^ mice when incubated in normal Ca^2+^- containing buffer. These responses mediated by external Ca^2+^ entry are most likely due to the well-established roles of proton-sensing ion channels, such as TRPV1 and acid-sensing ion channels (ASICs), in the activation of sensory neurons by acid (27). Furthermore, the preservation of these responses to capsaicin and extracellular Ca^2+^-mediated acid signaling in turn demonstrates that deletion of GPR68 had not significantly impaired the activation of other proton-stimulated receptors in sensory neurons.

Importantly, this pattern of impaired acid responsiveness translated to colonic afferents, which showed markedly reduced activation at pH 5.5 in the presence of the GPR68 antagonist Ogremorphin, and virtually no response to acid in tissue from GPR68^-/-^ mice across a pH range of 4.0-6.5. These findings reveal a previously unrecognized role for GPR68 in mediating acid-induced activation of sensory nerve endings in the colon, resonant with *ex vivo* recordings of esophageal afferents (47). Interestingly, this included proton concentrations (pH 4.0 – 6.0) sufficient to activate other known acid-sensing receptors known to contribute to acid-evoked stimulation of colonic afferents, such as TRPV1. Given that responses to capsaicin were unchanged in both colonic afferents and DRG neurons from GPR68^-/-^ mice, the near-complete loss of acid-evoked afferent signaling strongly suggests that GPR68 is essential for enabling or amplifying sensory neuron activation in response to acid, potentially through facilitation of other proton-sensitive pathways like TRPV1.

Further evidence supporting the role of GPR68 in acid-mediated activation of colonic nociceptors was provided by experiments using Ogerin, a positive allosteric modulator (PAM) that selectively shifts GPR68’s pH response curve (42) and enhances GPR68 signalling in various tissues (48-50). In line with this mechanism, Ogerin potentiated colonic afferent responses to acid at pH 6.0 and sensitised the colonic afferent response to luminal distension with weakly acidic (pH 6.8), but not neutral (pH 7.4), Krebs buffer. Sensitisation by Ogerin was observed only at noxious distension pressures, consistent with the selective expression of GPR68 in colonic nociceptors. This was further supported by single-unit recordings from mechanosensitive colonic afferent fibres, where Ogerin enhanced acid responses in units that responded exclusively to noxious levels of colorectal distension.

Despite modulating the activity of distension-sensitive fibres, and in contrast to recent reports suggesting that GPR68 can be activated by membrane stretch and shear stress (51, 52), consistent with the presence of a conserved Helix 8 structural motif shared among mechanosensitive GPCRs (53), we found no difference in the magnitude of the colonic afferent response to colorectal distension between wild-type and GPR68^-/-^ mice. While this finding does not exclude the possibility that GPR68 is mechanosensitive, it suggests a likely functional redundancy in its role in mechanotransduction within colonic afferents, especially when compared to the established contributions of other mechanosensitive channels such as TRPA1 and PIEZO2, whose loss results in a marked reduction in colonic afferent responses to distension (54, 55). Nevertheless, our data generated using Ogerin demonstrate that although GPR68 may not function as a primary mechanotransducer in colonic afferents, its activation can still enhance LSN sensitivity to mechanical stimuli.

In summary, our findings reveal a critical role of GPR68 in the activation of colonic afferents by acid and demonstrate its capacity to sensitize colonic afferent responses to noxious distension of the bowel, as shown through both pharmacological and genetic approaches.

These results position GPR68 as a key contributor to colonic nociception during tissue acidosis and support the need for further investigation into its role in pain signaling in GI disorders where acidosis is a prominent feature, such as colitis.

## Abbreviations

CD: Crohn’s disease
DRG: dorsal root ganglion
DSS: dextran sulphate sodium
ECS: extracellular solution
GI: gastrointestinal
GPCRs: G protein-coupled receptors
GPR68: G protein-coupled receptor 68
IBD: inflammatory bowel disease
LSN: lumbar splanchnic nerve
OGM: Ogremorphin
ONC: Ogerin negative control
TRPV1: transient receptor potential vanilloid 1
UC: ulcerative colitis

## Author Contributions

L.W.P. designed the research studies, conducted the experiments, acquired and analysed the data and wrote the manuscript. R.G., J.P.H., J.A.L. and A.R. acquired and analysed the data. T.P., S.N. and J.R.F.H analysed the data. T.R., G.R. and K.B. reviewed and revised the manuscript. F.W., E.St.J.S., and D.C.B. designed the research studies, obtained funding and wrote the manuscript.

## References

1. Fallingborg J, Christensen LA, Jacobsen BA, Rasmussen SN. Very low intraluminal colonic pH in patients with active ulcerative colitis. Dig Dis Sci. 1993;38(11):1989–93.

2. Nugent SG, Kumar D, Rampton DS, Evans DF. Intestinal luminal pH in inflammatory bowel disease: possible determinants and implications for therapy with aminosalicylates and other drugs. Gut. 2001;48(4):571–7.

3. Press AG, Hauptmann IA, Hauptmann L, Fuchs B, Fuchs M, Ewe K, et al. Gastrointestinal pH profiles in patients with inflammatory bowel disease. Aliment Pharmacol Ther. 1998;12(7):673-8.

4. Sasaki Y, Hada R, Nakajima H, Fukuda S, Munakata A. Improved localizing method of radiopill in measurement of entire gastrointestinal pH profiles: colonic luminal pH in normal subjects and patients with Crohn’s disease. Am J Gastroenterol. 1997;92(1):114–8.

5. Vernia P, Caprilli R, Latella G, Barbetti F, Magliocca FM, Cittadini M. Fecal lactate and ulcerative colitis. Gastroenterology. 1988;95(6):1564–8.

6. Caprilli R, Frieri G, Latella G, Vernia P, Santoro ML. Faecal excretion of bicarbonate in ulcerative colitis. Digestion. 1986;35(3):136–42.

7. Hausmann M, Seuwen K, de Valliere C, Busch M, Ruiz PA, Rogler G. Role of pH-sensing receptors in colitis. Pflugers Arch. 2024;476(4):611–22.

8. Lardner A. The effects of extracellular pH on immune function. J Leukoc Biol. 2001;69(4):522–30.

9. Chen X, Jaiswal A, Costliow Z, Herbst P, Creasey EA, Oshiro-Rapley N, et al. pH sensing controls tissue inflammation by modulating cellular metabolism and endo-lysosomal function of immune cells. Nat Immunol. 2022;23(7):1063–75.

10. Yu XW, Hu ZL, Ni M, Fang P, Zhang PW, Shu Q, et al. Acid-sensing ion channels promote the inflammation and migration of cultured rat microglia. Glia. 2015;63(3):483–96.

11. Frasch SC, McNamee EN, Kominsky D, Jedlicka P, Jakubzick C, Zemski Berry K, et al. G2A Signaling Dampens Colitic Inflammation via Production of IFN-gamma. J Immunol. 2016;197(4):1425–34.

12. Pattison LA, Rickman RH, Hilton H, Dannawi M, Wijesinghe SN, Ladds G, et al. Activation of the proton-sensing GPCR, GPR65 on fibroblast-like synoviocytes contributes to inflammatory joint pain. Proc Natl Acad Sci U S A. 2024;121(51):e2410653121.

13. Dai SP, Hsieh WS, Chen CH, Lu YH, Huang HS, Chang DM, et al. TDAG8 deficiency reduces satellite glial number and pro-inflammatory macrophage number to relieve rheumatoid arthritis disease severity and chronic pain. J Neuroinflammation. 2020;17(1):170.

14. Miltz W, Velcicky J, Dawson J, Littlewood-Evans A, Ludwig MG, Seuwen K, et al. Design and synthesis of potent and orally active GPR4 antagonists with modulatory effects on nociception, inflammation, and angiogenesis. Bioorg Med Chem. 2017;25(16):4512–25.

15. Osthues T, Zimmer B, Rimola V, Klann K, Schilling K, Mathoor P, et al. The Lipid Receptor G2A (GPR132) Mediates Macrophage Migration in Nerve Injury-Induced Neuropathic Pain. Cells. 2020;9(7).

16. Sisignano M, Fischer MJM, Geisslinger G. Proton-Sensing GPCRs in Health and Disease. Cells. 2021;10(8).

17. Ludwig MG, Vanek M, Guerini D, Gasser JA, Jones CE, Junker U, et al. Proton-sensing G-protein-coupled receptors. Nature. 2003;425(6953):93-8.

18. Matsingos C, Howell LA, McCormick PJ, Fornili A. Elucidating the Activation Mechanism of the Proton-sensing GPR68 Receptor. J Mol Biol. 2024;436(16):168688.

19. Guo L, Zhu K, Zhong YN, Gao M, Liu J, Qi Z, et al. Structural basis and biased signaling of proton sensation by GPCRs mediated by extracellular histidine rearrangement. Mol Cell. 2025;85(8):1658–73 e7.

20. Howard MK, Hoppe N, Huang XP, Mitrovic D, Billesbolle CB, Macdonald CB, et al. Molecular basis of proton sensing by G protein-coupled receptors. Cell. 2025;188(3):671–87 e20.

21. Hutter S, van Haaften WT, Hunerwadel A, Baebler K, Herfarth N, Raselli T, et al. Intestinal Activation of pH-Sensing Receptor OGR1 [GPR68] Contributes to Fibrogenesis. J Crohns Colitis. 2018;12(11):1348–58.

22. de Valliere C, Wang Y, Eloranta JJ, Vidal S, Clay I, Spalinger MR, et al. G Protein-coupled pH-sensing Receptor OGR1 Is a Regulator of Intestinal Inflammation. Inflamm Bowel Dis. 2015;21(6):1269–81.

23. de Valliere C, Babler K, Busenhart P, Schwarzfischer M, Maeyashiki C, Schuler C, et al. A Novel OGR1 (GPR68) Inhibitor Attenuates Inflammation in Murine Models of Colitis. Inflamm Intest Dis. 2021;6(3):140–53.

24. Mohebbi N, Benabbas C, Vidal S, Daryadel A, Bourgeois S, Velic A, et al. The proton-activated G protein coupled receptor OGR1 acutely regulates the activity of epithelial proton transport proteins. Cell Physiol Biochem. 2012;29(3-4):313–24.

25. de Valliere C, Cosin-Roger J, Baebler K, Schoepflin A, Mamie C, Mollet M, et al. pH-Sensing G Protein-Coupled Receptor OGR1 (GPR68) Expression and Activation Increases in Intestinal Inflammation and Fibrosis. Int J Mol Sci. 2022;23(3).

26. de Valliere C, Cosin-Roger J, Simmen S, Atrott K, Melhem H, Zeitz J, et al. Hypoxia Positively Regulates the Expression of pH-Sensing G-Protein-Coupled Receptor OGR1 (GPR68). Cell Mol Gastroenterol Hepatol. 2016;2(6):796–810.

27. Pattison LA, Callejo G, St John Smith E. Evolution of acid nociception: ion channels and receptors for detecting acid. Philos Trans R Soc Lond B Biol Sci. 2019;374(1785):20190291.

28. Khan Z, Jung M, Crow M, Mohindra R, Maiya V, Kaminker JS, et al. Whole genome sequencing across clinical trials identifies rare coding variants in GPR68 associated with chemotherapy-induced peripheral neuropathy. Genome Med. 2023;15(1):45.

29. Huang CW, Tzeng JN, Chen YJ, Tsai WF, Chen CC, Sun WH. Nociceptors of dorsal root ganglion express proton-sensing G-protein-coupled receptors. Mol Cell Neurosci. 2007;36(2):195–210.

30. Higham JP, Bhebhe CN, Gupta RA, Tranter MM, Barakat FM, Dogra H, et al. Transcriptomic profiling reveals a pronociceptive role for angiotensin II in inflammatory bowel disease. Pain. 2024;165(7):1592–604.

31. Hockley JRF, Taylor TS, Callejo G, Wilbrey AL, Gutteridge A, Bach K, et al. Single-cell RNAseq reveals seven classes of colonic sensory neuron. Gut. 2019;68(4):633–44.

32. Pattison LA, Cloake A, Chakrabarti S, Hilton H, Rickman RH, Higham JP, et al. Digging deeper into pain: an ethological behavior assay correlating well-being in mice with human pain experience. Pain. 2024;165(8):1761–73.

33. Taylor TS, Konda P, John SS, Bulmer DC, Hockley JRF, Smith ESJ. Galanin suppresses visceral afferent responses to noxious mechanical and inflammatory stimuli. Physiol Rep. 2020;8(2):e14326.

34. Robinson DR, McNaughton PA, Evans ML, Hicks GA. Characterization of the primary spinal afferent innervation of the mouse colon using retrograde labelling. Neurogastroenterol Motil. 2004;16(1):113–24.

35. Barker KH, Higham JP, Pattison LA, Taylor TS, Chessell IP, Welsh F, et al. Sensitization of colonic nociceptors by TNFalpha is dependent on TNFR1 expression and p38 MAPK activity. J Physiol. 2022;600(16):3819–36.

36. Gupta RA, Higham JP, Pearce A, Urriola-Munoz P, Barker KH, Paine L, et al. GPR35 agonists inhibit TRPA1-mediated colonic nociception through suppression of substance P release. Pain. 2025;166(3):596–613.

37. Meng MY, Paine LW, Sagnat D, Bello I, Oldroyd S, Javid F, et al. TRPV4 stimulates colonic afferents through mucosal release of ATP and glutamate. Br J Pharmacol. 2025;182(6):1324–40.

38. Hockley JR, Tranter MM, McGuire C, Boundouki G, Cibert-Goton V, Thaha MA, et al. P2Y Receptors Sensitize Mouse and Human Colonic Nociceptors. J Neurosci. 2016;36(8):2364–76.

39. Ness TJ, Gebhart GF. Colorectal distension as a noxious visceral stimulus: physiologic and pharmacologic characterization of pseudaffective reflexes in the rat. Brain Res. 1988;450(1-2):153–69.

40. Hughes PA, Brierley SM, Martin CM, Brookes SJ, Linden DR, Blackshaw LA. Post-inflammatory colonic afferent sensitisation: different subtypes, different pathways and different time courses. Gut. 2009;58(10):1333–41.

41. Jordt SE, Tominaga M, Julius D. Acid potentiation of the capsaicin receptor determined by a key extracellular site. Proc Natl Acad Sci U S A. 2000;97(14):8134–9.

42. Huang XP, Karpiak J, Kroeze WK, Zhu H, Chen X, Moy SS, et al. Allosteric ligands for the pharmacologically dark receptors GPR68 and GPR65. Nature. 2015;527(7579):477-83.

43. Bielefeldt K, Davis B, Binion DG. Pain and inflammatory bowel disease. Inflamm Bowel Dis. 2009;15(5):778–88.

44. Sato K, Mogi C, Mighell AJ, Okajima F. A missense mutation of Leu74Pro of OGR1 found in familial amelogenesis imperfecta actually causes the loss of the pH-sensing mechanism. Biochem Biophys Res Commun. 2020;526(4):920–6.

45. Saxena H, Deshpande DA, Tiegs BC, Yan H, Battafarano RJ, Burrows WM, et al. The GPCR OGR1 (GPR68) mediates diverse signalling and contraction of airway smooth muscle in response to small reductions in extracellular pH. Br J Pharmacol. 2012;166(3):981–90.

46. Ichimonji I, Tomura H, Mogi C, Sato K, Aoki H, Hisada T, et al. Extracellular acidification stimulates IL-6 production and Ca(2+) mobilization through proton-sensing OGR1 receptors in human airway smooth muscle cells. Am J Physiol Lung Cell Mol Physiol. 2010;299(4):L567–77.

47. Ru F, Banovcin P, Jr., Kollarik M. Acid sensitivity of the spinal dorsal root ganglia C-fiber nociceptors innervating the guinea pig esophagus. Neurogastroenterol Motil. 2015;27(6):865–74.

48. Bell TJ, Nagel DJ, Woeller CF, Kottmann RM. Ogerin mediated inhibition of TGF-beta(1) induced myofibroblast differentiation is potentiated by acidic pH. PLoS One. 2022;17(7):e0271608.

49. Li X, Xia K, Zhong C, Chen X, Yang F, Chen L, et al. Neuroprotective effects of GPR68 against cerebral ischemia-reperfusion injury via the NF-kappaB/Hif-1alpha pathway. Brain Res Bull. 2024;216:111050.

50. Karki P, Ke Y, Zhang C, Promnares K, Li Y, Williams CH, et al. GPR68 Mediates Lung Endothelial Dysfunction Caused by Bacterial Inflammation and Tissue Acidification. Cells. 2024;13(24).

51. Xu J, Mathur J, Vessieres E, Hammack S, Nonomura K, Favre J, et al. GPR68 Senses Flow and Is Essential for Vascular Physiology. Cell. 2018;173(3):762–75 e16.

52. Wei WC, Bianchi F, Wang YK, Tang MJ, Ye H, Glitsch MD. Coincidence Detection of Membrane Stretch and Extracellular pH by the Proton-Sensing Receptor OGR1 (GPR68). Curr Biol. 2018;28(23):3815–23 e4.

53. Erdogmus S, Storch U, Danner L, Becker J, Winter M, Ziegler N, et al. Helix 8 is the essential structural motif of mechanosensitive GPCRs. Nat Commun. 2019;10(1):5784.

54. Xie Z, Feng J, Hibberd TJ, Chen BN, Zhao Y, Zang K, et al. Piezo2 channels expressed by colon-innervating TRPV1-lineage neurons mediate visceral mechanical hypersensitivity. Neuron. 2023;111(4):526–38 e4.

55. Cattaruzza F, Spreadbury I, Miranda-Morales M, Grady EF, Vanner S, Bunnett NW. Transient receptor potential ankyrin-1 has a major role in mediating visceral pain in mice. Am J Physiol Gastrointest Liver Physiol. 2010;298(1):G81–91.

