## Supplemental Figures for "GPR68, a proton-sensing GPCR, mediates acid-induced visceral nociception"

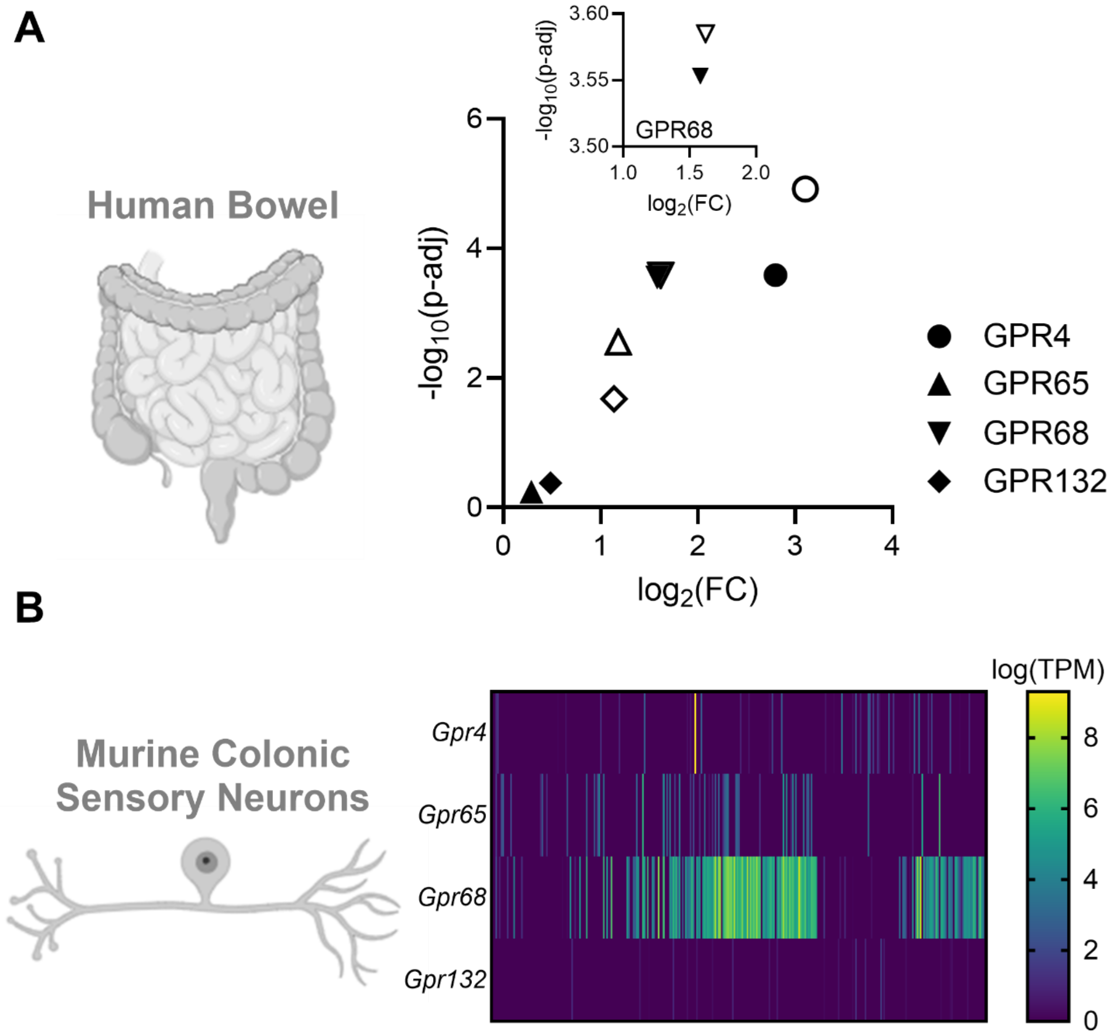


**Supplemental Figure 1. Expression of four proton-sensing GPCRs in UC and CD and expression in colon-innervating sensory neurons.** (A) Expression of selected proton-sensing GPCRs in UC (filled symbols) and CDN (open symbols) given as log2 fold change compared to non-inflamed controls with the associated FDR-adjusted p-value (as -log10). Inset: expanded view highlighting *GPR68* expression (26). (B) Heatmap showing the expression (logTPM) of *Gpr4*, *Gpr65*, *Gpr68* and *Gpr132* in murine colonic sensory neurons (data redrawn from Hockley et al. (27)).

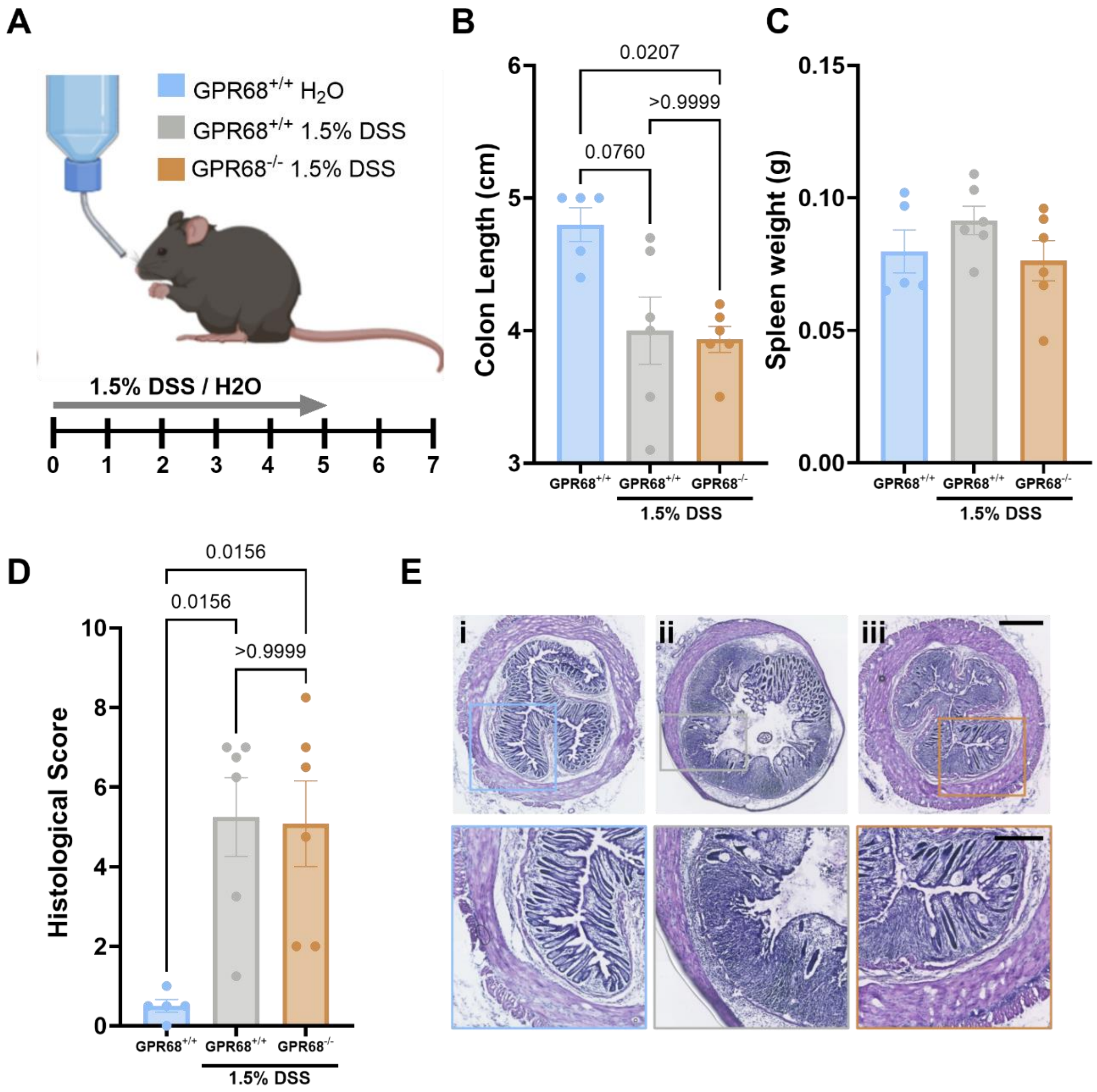


**Supplemental Figure 2. Additional parameters of the DSS-induced colitis model in GPR68$^{+/+}$ and GPR68$^{-/-}$ mice.** (A) Schematic representation of the DSS model and experimental timeline. (B) colon length and spleen weight (C) measured on day 7 following administration of H2O or 1.5% DSS in GPR68+/+ and GPR68-/- mice (colon length: H2O control vs GPR68+/+ DSS: P = 0.0760; H2O control vs GPR68-/- DSS: P = 0.0207; GPR68+/+ DSS vs GPR68-/- DSS: P = >0.9999; Kruskal-Wallis test (main effect: P = 0.01; H = 8.129) with Dunn's multiple comparison (Z = 2.236, 2.702, 0.4886, respectively)) (spleen weight: H2O control vs GPR68+/+ DSS: P = 0.6834; H2O control vs GPR68-/- DSS: P = >0.9999; GPR68+/+ DSS vs GPR68-/- DSS: P = 0.4333; Kruskal-Wallis test (main effect: P = 0.3045; H = 2.467) with Dunn's multiple comparison (Z = 1.206, 0.1855, 0.1.460, respectively)). (D) Histological scores of colon sections based on inflammation, crypt damage and ulceration (H2O control vs GPR68+/+ DSS: P = 0.0156; H2O control vs GPR68-/- DSS: P = 0.0156; GPR68+/+ DSS vs GPR68-/- DSS: P = >0.9999; Kruskal-Wallis test (main effect: P = 0.0019; H = 10.11) with Dunn's multiple comparison (Z = 2.795, 2.795, 0.000, respectively)). (G) Representative H&E staining of colon sections across experimental groups (i) H2O control (ii) GPR68+/+ 1.5% DSS and (iii) GPR68-/- 1.5% DSS. Scale bars: full cross-sections = 500 µm, higher magnification insets = 250 µm. (N = 5-6 animals).

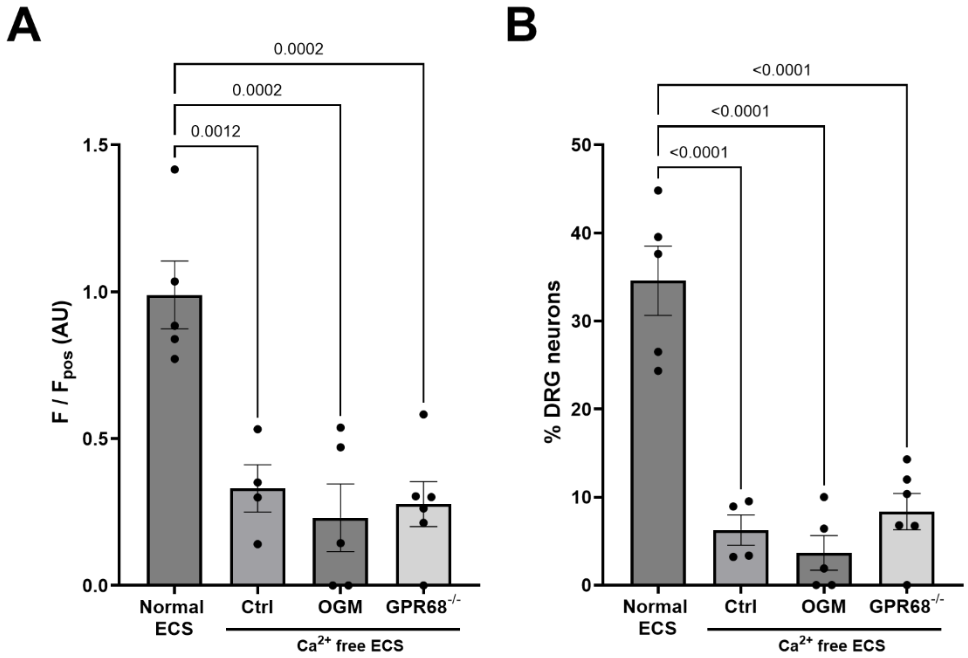


**Supplemental Figure 3. Attenuation of capsaicin response in DRG neurons following extracellular $Ca^{2+}$ removal.** (A) Peak responses and (B) the proportion of DRG neurons responding to capsaicin (1 μM) in normal ECS (data taken from Supplemental Figure 1) and $Ca^{2+}$-free ECS across treatment groups (Peak: Normal ECS vs $Ca^{2+}$-free ECS, Control (P = 0.0012), vs OGM (P = 0.0002), vs $GPR68^{-/-}$ (P = 0.0002), one-way ANOVA (main effect: P = 0.0001, F(3, 16) = 13.11) with Dunnett's multiple comparisons test; n = 4-6 dishes from N = 3-5 animals) (Proportion of DRG neurons: Normal ECS vs $Ca^{2+}$-free ECS, Control (P = <0.0001), vs OGM (P = <0.0001), vs $GPR68^{-/-}$ (P = <0.0001), one-way ANOVA (main effect: P = <0.0001, F(3, 16) = 29.79) with Dunnett's multiple comparisons test; n = 4-6 dishes from N = 3-5 animals).

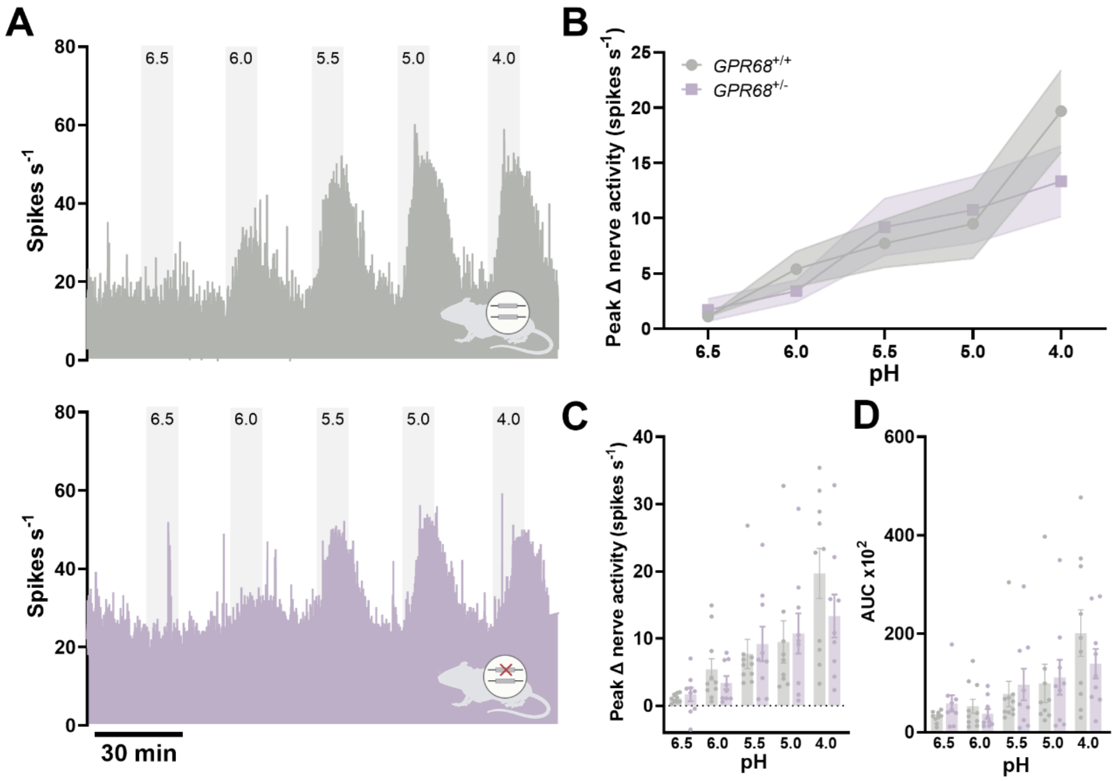


**Supplemental Figure 4. Comparable colonic afferent responses to acid in GPR68$^{+/+}$ and GPR68$^{+/-}$ mice.** (A) Representative rate histograms and (B) mean response profiles of GPR68$^{+/+}$ (grey) and GPR68$^{+/-}$ (lilac) colonic afferent activity in response to pH 6.5-4.0; shaded area is SEM. (GPR68$^{+/+}$ vs GPR68$^{+/-}$ main effect: P = 0.5185; two-way ANOVA; F(1, 83) = 0.4204). (C) Peak afferent activity at each pH (GPR68$^{+/+}$ vs GPR68$^{+/-}$ unpaired t-test (or non-parametric equivalent); pH 6.5, P = 0.5770, t, df = 0.5694, 16; pH 6.0, P = 0.2428 (U = 30); pH 5.5, P = 0.7197 (U = 40); pH 5.0, P = 0.6665 (U = 35); pH 4.0, P = 0.2179, t, df = 1.279, 17; N = 9-10 animals). (D) Area under the curve (AUC) for responses across the pH range (GPR68$^{+/+}$ vs GPR68$^{+/-}$ unpaired t-test (or non-parametric equivalent); pH 6.5, P = 0.0939 (U = 21); pH 6.0, P = 0.4470 (U = 35); pH 5.5, P = 0.9682 (U = 44); pH 5.0, P = 0.8633 (U = 38); pH 4.0, P = 0.4002 (U = 34); N = 9-10 animals). Wild-type data shown here are the same as those presented in Figure 4 and are included for direct comparison with GPR68$^{+/-}$ mice.

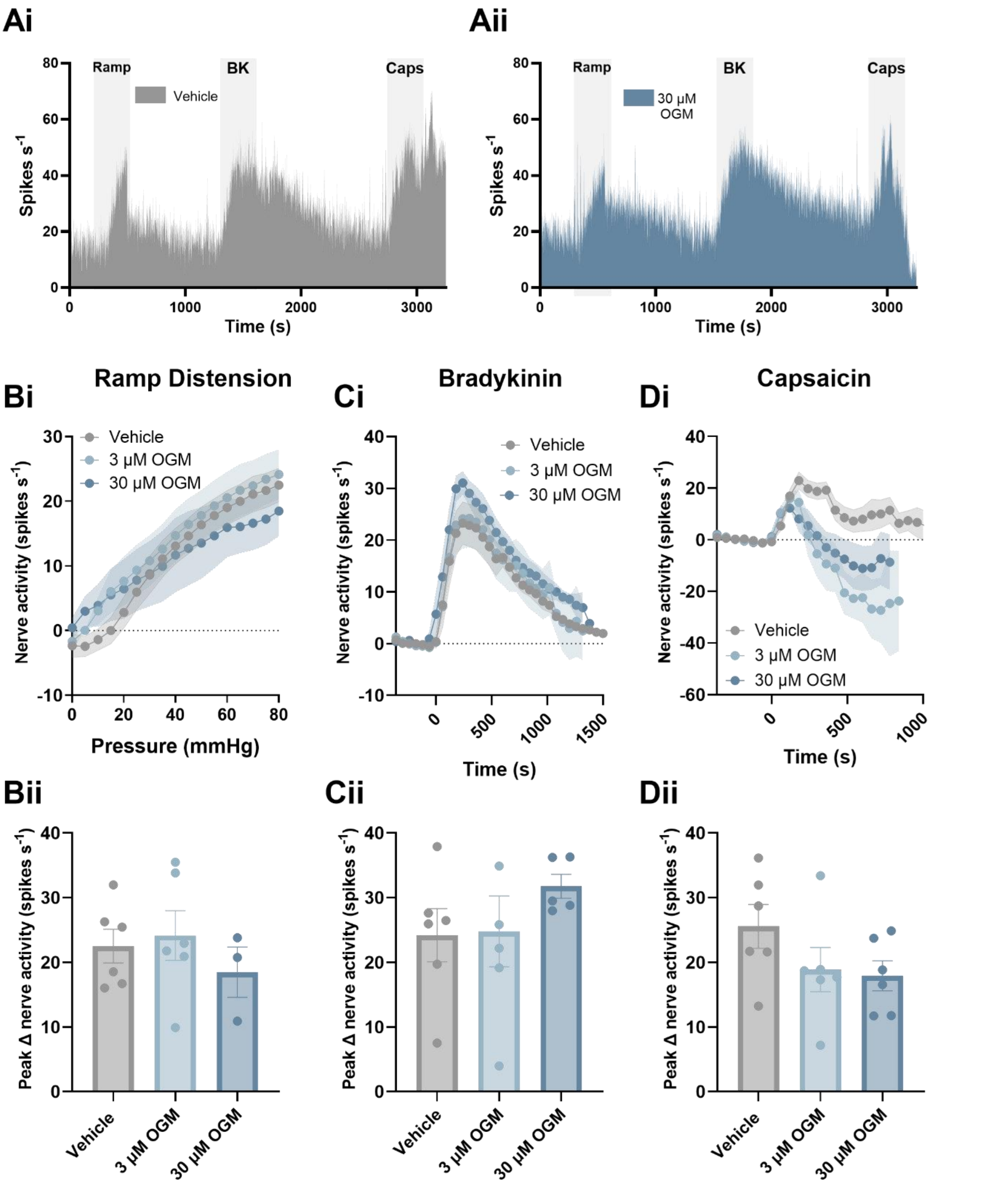


**Supplemental Figure 5. Comparable colonic afferent responses to mechanical and chemical (bradykinin and capsaicin) stimuli following treatment with OGM.** (A) Representative rate histogram of colonic afferent responses to ramp distension (Ramp), bradykinin (BK; 1 μM) and capsaicin (Caps; 1 μM) following treatment with 3 μM or 30 μM OGM (GPR68 antagonist) or vehicle. (Bi) Afferent response profile (P = 0.7647; two-way ANOVA; N = 3-6) and (Bii) peak change in afferent activity at 80 mmHg in response to ramp distension (vehicle vs 3 μM OGM, P = 0.9135; vehicle vs 30 μM OGM, P = 0.7072; one-way ANOVA (main effect: P = 0.6056; $F(2, 12) = 0.5232$) with Dunnett's multiple comparison test; N = 3-6). (Ci) Response profile (P = 0.4615; two-way ANOVA; N = 5-6) and (Cii) peak change in afferent activity in response to bradykinin (1 μM) stimulation (vehicle vs 3 μM OGM, $P = 0.9945$; vehicle vs 30 μM OGM, $P = 0.4618$; one-way ANOVA (main effect: $P = 0.4315$; $F(2, 14) = 0.8931$) with Dunnett's multiple comparison test; N = 5-6). (Di) Response profile ($P = 0.0524$; two-way ANOVA; N = 6) and (Dii) peak change in afferent activity in response to capsaicin (1 μM) stimulation (vehicle vs 3 μM OGM, $P = 0.3043$; vehicle vs 30 μM OGM, $P = 0.2195$; one-way ANOVA (main effect: $P = 0.1959$; $F(2, 15) = 1.821$) with Dunnett's multiple comparison test; N = 6).
